# Root phenotypic plasticity improves yield stability when directed toward an adaptive integrated phenotype

**DOI:** 10.64898/2026.08.10.744026

**Authors:** Ivan Lopez-Valdivia, Akshay B. Tawale, Matías Schierenbeck, Daniele Sandoni, Dylan H. Jones, Gwendolyn K. Kirschner, Hannah M. Schneider

## Abstract

Root phenotypic plasticity is often proposed to improve crop performance under stress, yet it remains unclear how much plasticity is beneficial and whether adaptive responses require changes across many traits or adjustments in few specific traits. Using public data of 6,500 field-grown maize and barley plants, this study examined the extent and distribution of root plasticity, and when it is associated with yield stability.

We quantified root plasticity across nine anatomical and architectural traits using complementary statistical models and applied a feature-discovery framework to identify the drought-associated optimal integrated phenotypes and determine whether plasticity toward these phenotypes improved yield stability.

More plasticity did not mean greater yield stability. Neither the number of plastic traits nor the magnitude of plastic responses predicted yield stability. Rather, we identified species-specific high-yielding, stable integrated phenotypes defined by distinct trait configurations. Critically, genotypes whose plastic responses moved their root phenotype toward these targets achieved greater yield stability, whereas movement away from them was associated with lower stability.

Root plasticity is adaptive when it shifts root phenotypes towards an optimal integrated phenotype. These findings show that the value of plasticity depends on the trajectory of phenotypic change rather than its magnitude alone.

## Introduction

Global agriculture faces increasing challenges from drought, diminishing freshwater resources, shifting precipitation patterns, ongoing soil degradation, and rising production costs (Thingujam et al., 2025; Tebaldi and Lobell, 2008). Among these factors, drought remains one of the greatest threats to crop productivity and food security because it directly limits plant biomass accumulation, reproductive development, thereby disrupting source– sink relationships that determine grain number and grain filling in cereals (Mondal et al., 2026; Li et al., 2026). As climate variability increases, improving yield stability across contrasting environments has become an important objective for crop improvement programs.

Yield stability is broadly defined as the ability of a genotype to maintain consistent performance despite environmental variation (Knapp and van der Heijden, 2018; Sadras et al., 2026). For farmers operating in unpredictable environments, stable yields are often as important as maximizing yield potential (Tollenaar and Lee, 2002). Achieving stable production requires plants to maintain resource acquisition and reproductive development under fluctuating environmental conditions. Because roots determine where and how effectively plants acquire soil water and nutrients, root phenotypes are widely recognized as important contributors to drought adaptation and yield stability (Strock et al., 2026; Lynch and Wojciechowski 2015; Siangliw et al., 2022; Lopez-Valdivia et al., 2025). Phenotypic plasticity describes the capacity of an organism to alter its phenotype in response to varying environmental conditions (Bradshaw 1965, Sultan 1995). In plants, plastic responses can occur through changes in physiology, morphology, anatomy, development, or resource allocation (Wang et al., 2026). Root phenotypic plasticity specifically refers to environmentally induced changes in root architecture, anatomy, and function that alter how roots explore the soil and acquire resources (Schneider et al., 2020a, 2020b; Schneider and Lynch, 2020). Phenotypic plasticity can be adaptive (beneficial), maladaptive (detrimental), or neutral with respect to its impact on plant fitness (Schneider, 2022; Sadras et al., 2026; King and Hadfield, 2019). Because soil resources are highly heterogeneous in space and time, root plasticity has frequently been proposed as an adaptive mechanism that enables plants to maintain performance under environmental stress (Schneider and Lynch 2020; Karlova et al., 2021).

Numerous root traits exhibit plastic responses to drought. In response to varying environments, a wide range of crop species exhibit root plasticity, including changes in root angle, lateral root branching density, lateral root length, rooting depth, cortical anatomy, and xylem vessel size, and number (Ehdaie et al., 2012; Wasson et al., 2012; Zhan et al., 2015; Gao and Lynch, 2016; Hazman and Brown, 2018; Koehler et al 2025). Plastic responses such as hydropatterning can direct lateral root development toward water-rich microsites (Bao et al., 2014; Giehl and von Wirén, 2018), while modifications of root anatomy can alter the metabolic costs and hydraulic properties of root systems (Prince et al., 2017; Kadam et al., 2017). Several studies have linked individual plastic responses to improved plant performance under drought. For example, plasticity in root architectural traits has been associated with yield stability in rice (*Oryza sativa* L.) (Sandhu et al., 2016; Xie et al., 2021), while plastic changes in root anatomy have been associated with improved water acquisition and stress tolerance in rice, wheat (*Triticum aestivum* L.), and soybean (*Glycine max* (L.)) (Niones et al., 2012, 2013; Prince et al., 2017; Kadam et al., 2017). Despite extensive evidence that individual root traits respond plastically to drought, the relationship between root plasticity and yield stability remains poorly understood. Most studies have focused on individual traits in isolation, making it difficult to determine whether plasticity is beneficial or whether only specific plastic responses contribute to improved performance. Root systems are complex and composed of many interacting architectural and anatomical traits, each capable of responding independently to environmental conditions. Increasing evidence indicates that drought adaptation emerges from coordinated interactions among root anatomical and architectural traits rather than from individual traits acting independently. For example, Klein et al. (2020) demonstrated that multiple integrated root phenotypes were associated with improved drought tolerance in maize (*Zea mays* L.) and Ajmera et al. (2022) proposed integrated root phenotypes for rice associated with improved nitrogen capture, suggesting that adaptive value resides in specific phenotypic configurations rather than isolated trait states. Given the importance of specific phenotypic configurations for plant adaptation, it is plausible that the adaptive value of plasticity may not depend on the total amount of phenotypic change expressed by the root system, but rather on whether those changes move the plant toward a more favorable integrated phenotype.

Recent studies support this possibility. While root plasticity is often assumed to improve environmental adaptation, phenotypic plasticity frequently shows weak relationships with yield stability (Xie et al., 2021; Sadras et al., 2026). These observations suggest that adaptive plasticity may depend less on the magnitude of phenotypic change and more on the direction of that change relative to an optimal phenotype. Under this framework, the same plastic response may be beneficial in one genotype but neutral or even detrimental in another depending on the constitutive phenotype from which the response occurs.

We hypothesized that indiscriminate root plasticity does not improve yield or yield stability but that the benefit of plasticity comes from movement towards a beneficial target. Rather than evaluating plasticity as a generalized whole-system response, we propose viewing it as a directed trajectory through phenotype space. To map these trajectories, we implemented a data-driven feature discovery pipeline on published root anatomical and architectural datasets from 429 maize and 192 barley (*Hordeum vulgare* L.) accessions grown under drought and well-watered field conditions. Specifically, we asked whether (i) the presence of plasticity across multiple root traits is associated with yield stability, (ii) the cumulative magnitude of root plasticity influences yield stability, (iii) plasticity in specific root traits contributes to yield stability, and (iv) a mathematical framework can identify an optimal integrated root phenotype and quantify the adaptive value of plasticity moving toward it. By combining comparative analyses across two crop species, we aim to determine in what instances root plasticity contributes to yield stability and whether adaptive value depends on movement toward favorable integrated phenotypes.

## Materials and Methods

### Data collection

Publicly available root phenotypic datasets were obtained for two crop species grown under field conditions with contrasting water availability. The datasets included 429 modern inbred maize accessions grown in Arizona, USA during 2015 and 2016 (Schneider et al., 2020a; Schneider et al., 2020b) and 192 spring barley accessions (combination of 2- and 6-row barley of old cultivars, landraces, and modern breeding materials) grown in Bonn, Germany during 2013, 2014, and 2015 (Oyiga et al., 2020). The drought treatments differed among species, and each case were designed to impose water limitation during critical developmental stages (drip irrigated and non-irrigated conditions for maize and rain-out shelter and rainfed conditions for barley). Maize was subjected to terminal drought beginning four weeks after planting. Barley experienced intermittent drought beginning at tiller initiation (BBCH21) and continuing until heading (BBCH51). To compare drought severity among studies, yield under drought was expressed relative to the mean yield under well-watered conditions. Yield is defined here as grams of grain per plant. Although crop species show different critical periods depending on the timing of stress (Carrera et al., 2024), the average yield reductions were similar in maize (34%) and barley (25%) (Figure S1). Further information about agronomic management is described in Schneider et al., (2020a), (2020b) and Oyiga et al., (2020).

### Root phenotypes

Root architectural traits were quantified using shovelomics (Trachsel et al., 2011), and root anatomical traits were quantified using laser ablation tomography (Hall et al., 2019). Nine common root phenotypes were available for maize and barley: root growth angle, lateral root branching frequency, lateral root length, root cross-sectional area, root cortical aerenchyma, cortical cell size, cortical cell file number, metaxylem vessel number, and median metaxylem vessel area (Table 1). The description of these traits and how they were collected is described in Schneider et al., 2020a; Schneider et al., 2020b; Oyiga et al., 2020. Flowering time expressed in growing degree days was available for maize, but not for barley.

**Table 1.** Root anatomical and architectural traits used for this study. Traits are described in Schneider et al., 2020a, b; Oyiga et al., 2020.

| Abbreviation | Trait |
| --- | --- |
| **CCFN** | Cortical cell file number (count) |
| **CCS** | Cortical cell size ($\mu\text{m}^2$) |
| **LL** | Lateral root length (mm) |
| **LRBF** | Lateral root branching frequency (branches per cm) |
| **RCA** | Root cortical aerenchyma (%) |
| **MXA** | Median area of metaxylem vessels ($\text{mm}^2$) |
| **NMXV** | Number of metaxylem vessels (count) |
| **RA** | Root angle (degrees) |
| **RXSA** | Root cross-sectional area ($\text{mm}^2$) |

### Data normalization

All analyses were performed in R version 4.5.2 (R Core Team) using RStudio version 2026.01.0+392. To facilitate comparisons among traits and species with different measurement scales, phenotypic data were normalized with the *dplyr* package (Wickham et al., 2023) using min–max scaling. Normalized trait values therefore ranged from 0 to 1 (Figures S2 and S3).

### Quantification of Plasticity

Phenotypic plasticity was quantified using three complementary approaches: Relative Distance Plasticity Index (RDPI), genotype-specific reaction norm slopes estimated a Finlay- Wilkinson Mixed Model (MM), and genotype-by-environment interaction scores derived from the Additive Main effects and Multiplicative Interaction (AMMI) analysis as described in Arenas et al., 2025.

## Relative Distance Plasticity Index (RDPI)

Plasticity magnitude and direction was quantified using the RDPI, calculated as:

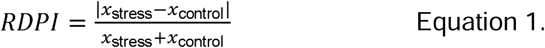

For each genotype, RDPI was calculated using trait values under drought and well-watered conditions. Both absolute RDPI values and directional RDPI values (without the absolute operator) were calculated. Absolute RDPI quantified the magnitude of plasticity, whereas directional RDPI preserved whether trait values increased or decreased in response to drought. The use of absolute and directional RDPI values are specified throughout the manuscript.

## Finlay Wilkinson Mixed Model (MM)

We calculated plasticity through a Finlay–Wilkinson mixed model (MM) using the *lme4* (Bates et al., 2015) and *emmeans* (Lenth, 2024) packages. We calculated environmental indices as the mean of the normalized trait across all genotypes within each unique environment-year-treatment combination. We then fitted a linear mixed model where the trait was modeled as a function of the genotype-by-environmental index interaction, with the environment ID (unique combination for year and treatment) as a random effect. Plasticity was defined as the genotype-specific slope (regression coefficient) of the trait response across the environmental indices.

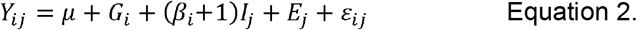

Where represents the observed trait value for genotype *i* in environment *j*; is the grand mean; is the fixed effect of the *i*-th genotype; is the continuous environmental index; is the genotype-specific slope representing phenotypic plasticity; represents the random effect of environment ID; and is the residual error. Plasticity slopes ( ) were considered statistically significant if their 95% confidence intervals did not overlap zero.

## Additive Main effects and Multiplicative Interaction (AMMI)

Genotype-by-environment interactions were evaluated using the AMMI model implemented in the *agricolae* package (Mendiburu, 2023). We extracted the scores from the first principal component (PC1) of the interaction and considered as the measure of plasticity. The formula for the AMMI model is expressed as:

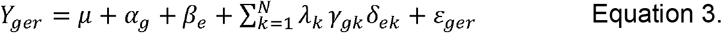

Where and represent genotype and environmental main effects, respectively; is the singular value of the *k*-th interaction principal component axis (IPCA); and and are the respective genotype and environmental eigenvectors for axis *k*. Genotype coordinates derived from the first principal component (PC1) were extracted.

To quantify plasticity at the scale of the entire root system, we developed two complementary metrics using nine root anatomical and architectural traits measured in maize and barley: Plastic Trait Count and Root System RDPI.

### Plastic Trait Count

The first metric quantified the number of root traits that exhibited significant plasticity in response to drought. For each genotype, individual traits were classified as plastic and given a score of ‘1’ or non-plastic and given a score of ‘0’ based on the plasticity criteria defined for RDPI, Finlay–Wilkinson mixed models, and AMMI analyses. To evaluate the presence or absence of root plasticity across genotypes, three distinct statistical criteria were applied depending on the metric used. For the Relative Distance Plasticity Index (RDPI), a trait- specific threshold was set at the 25^th^ percentile (Q1), genotypes scoring above this threshold were classified as plastic (1), while the bottom 25% least plastic genotypes were classified as non-plastic (0). For the Finlay-Wilkinson (MM) model, classification was strictly based on statistical significance (confidence intervals did not overlap zero), where genotypes with significant regression slopes were coded as plastic (1) and non-significant ones as non- plastic (0). Finally, for the AMMI model, genotypes were categorized as plastic (1) if their interaction scores (PC1) deviated from 0 by more than one standard deviation, and non- plastic (0) if they remained within this threshold. Scores were then summed across all nine traits to obtain a plastic trait count:

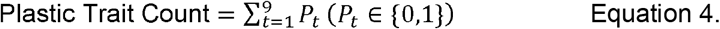

Where 0,1 indicates the presence (1) or absence (0) of plasticity for each trait t. This metric quantified the number of root phenotypes that responded plastically to drought for each genotype.

### Root system RDPI

The second metric quantified the cumulative magnitude of plasticity across the root system. For each genotype, absolute RDPI values were calculated for all nine root traits and summed:

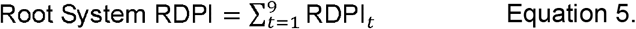

Unlike the plastic trait count, which only records whether a trait was plastic, root system RDPI is calculated as the sum of the magnitudes of the phenotypic responses across all traits *t*, providing a measure of the total magnitude of plasticity across a root system.

### Calculation of yield stability

For the purposes of this study, we define yield stability as reduced yield plasticity across environments. Because the metrics for plasticity operate on different scales, we applied model-specific transformations to calculate both trait stability and yield stability from their respective plasticity scores (Figure S4). For the Relative Distance Plasticity Index (RDPI), which produces positive values bounded between 0 and 1, stability was calculated by multiplying the original metric by -1. Conversely, because plasticity estimates derived from Mixed Models (MM) can take positive and negative values depending on the direction of the interaction, stability was calculated as the negative absolute value of the metric to capture the total magnitude of plasticity regardless of direction. For the Additive Main Effects and Multiplicative Interaction (AMMI) models, PC1 showed a positive correlation with the yield stability derived from RDPI (Figure S4), so we directly considered PC1 as a measured of yield stability. Under this unified framework, increasingly positive values represents greater stability, while increasingly negative values indicate greater plasticity and, consequently, lower stability.

### Bulk segregant analysis

To identify integrated phenotypes associated with yield stability, maize and barley genotypes were first classified into yield-stable and yield-unstable groups based on yield stability metrics (Figure S5). Within each group, root plasticity profiles using nine root traits were analyzed using Partition Around Medoids (PAM) clustering implemented in the *cluster* package (Maechler et al., 2019). Euclidean dissimilarity matrices were constructed using the daisy function, and the optimal number of clusters was determined using the *factoextra* (Kassambara and Mundt, 2020) and *NbClust* packages (Charrad et al., 2014). To improve cluster reliability, only genotypes showing strong membership in their assigned clusters (silhouette widths > 0.25) were retained for downstream analyses. In each species, two clusters were found for yield-stable and yield-unstable, they were classified as stable C1, stable C2, unstable C1 and unstable C2.

### Identification of drought-associated integrated phenotypes

To identify phenotypic combinations associated with drought performance, we employed a phenotype-space framework adapted from Klein et al. (2020) and Ajmera et al (2022). All possible combinations of root phenotypes were evaluated (Table S1, S2 and S3), ranging from two traits to the full trait set (>1,500 unique combinations). For each trait combination, PAM clustering was performed across cluster numbers ranging from 2 to 6 (Table S4 and S5). Cluster quality was evaluated using average silhouette width, and only cluster configurations with mean silhouette values greater than 0.25 were retained.

To determine whether a given phenotypic configuration captured biologically meaningful variation, yield under drought was compared among clusters using analysis of variance (ANOVA) followed by Tukey’s Honest Significant Difference (HSD) tests implemented in the *agricolae* package (Mendiburu, 2023). The trait combination and cluster configuration that produced the strongest separation of drought yield among clusters were retained as the candidate drought-associated integrated phenotype (Table S6 and S7). The optimum phenotype was considered as the one with the highest cluster structure and the most significant P value for yield differences under drought (Table 2 and Table 3). However, several other phenotypic combinations showed strong structure and yield differences under drought (Table S6 and S7).

**Table 2:**
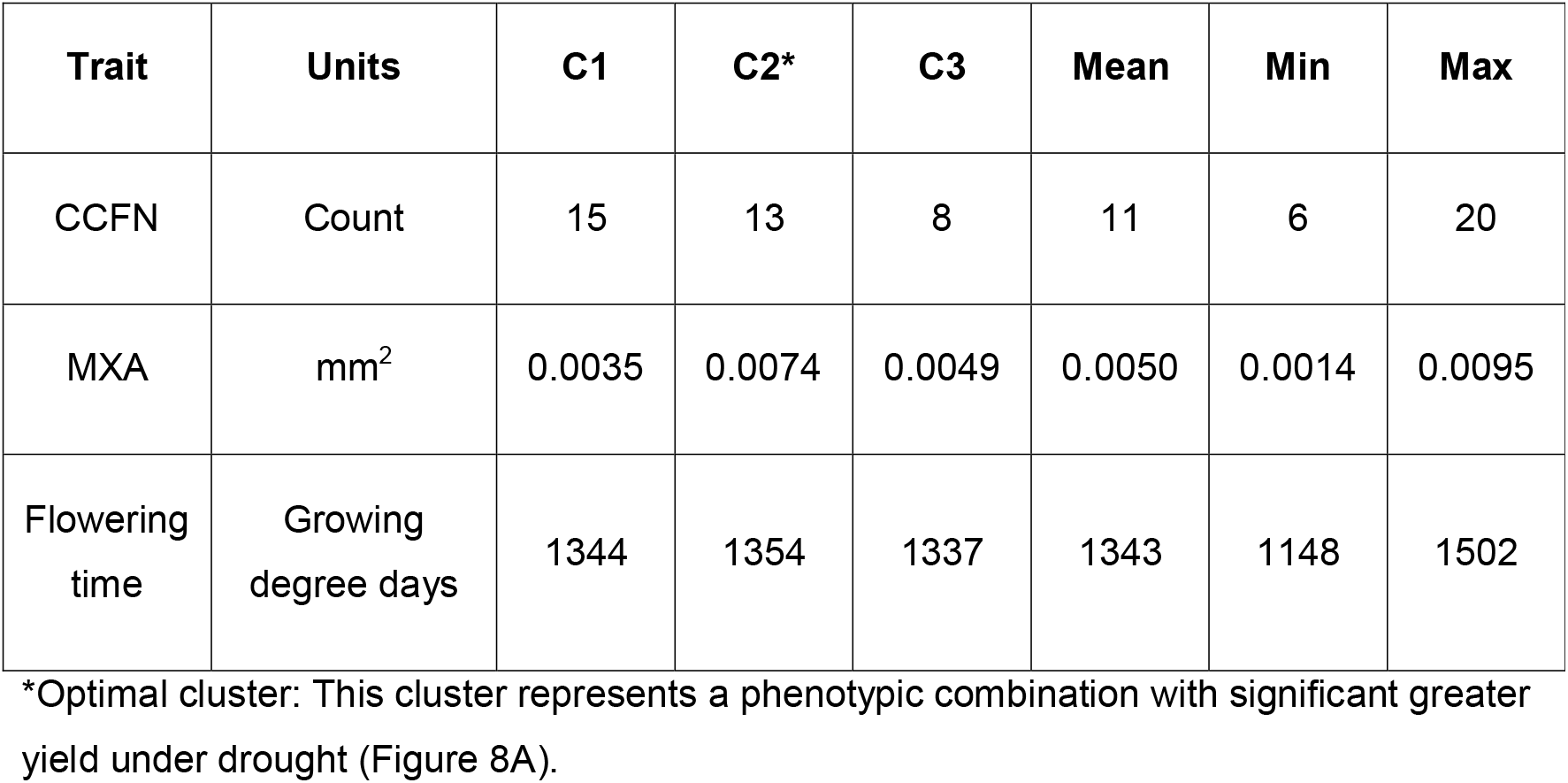
Average root phenotypes of clusters 1 (C1), 2 (C2) and 3 (C3) in maize. Cluster 2 shows higher yield and yield stability compared to clusters 1 and 3. The mean, minimum (Min), and maximum (Max) columns represent the metrics calculated across all maize genotypes under drought stress.

| Trait | Units | C1 | C2* | C3 | Mean | Min | Max |
| --- | --- | --- | --- | --- | --- | --- | --- |
| CCFN | Count | 15 | 13 | 8 | 11 | 6 | 20 |
| MXA | mm<sup>2</sup> | 0.0035 | 0.0074 | 0.0049 | 0.0050 | 0.0014 | 0.0095 |
| Flowering time | Growing degree days | 1344 | 1354 | 1337 | 1343 | 1148 | 1502 |

\*Optimal cluster: This cluster represents a phenotypic combination with significant greater yield under drought (Figure 8A).

**Table 3:** Average root phenotypes of cluster 1 (C1), 2 (C2), and 3 (C3) in barley. Cluster 1 shows higher yield and yield stability compared to clusters 2 and 3. The mean, minimum (Min), and maximum (Max) columns represent the metrics calculated across all barley genotypes under drought stress.

| Trait | Units | C1* | C2 | C3 | Mean | Min | Max |
| --- | --- | --- | --- | --- | --- | --- | --- |
| LRBF | Branches per cm | 7.32 | 7.69 | 6.75 | 7.10 | 5.57 | 8.50 |
| RXSA | mm<sup>2</sup> | 1.35 | 0.93 | 0.63 | 0.97 | 0.49 | 1.66 |
| CCS | μm<sup>2</sup> | 162.88 | 162.53 | 162.03 | 162.45 | 154 | 177.00 |

\*Optimal cluster: This cluster represents a phenotypic combination with significant greater yield under drought (Figure 8B).

### Definition of a drought-associated integrated phenotype

The cluster exhibiting the highest mean yield under drought was designated as the optimum phenotype. The centroid of this cluster was calculated as the multivariate mean of all traits included in the selected phenotype combination (Table 2 and 3). This centroid served as the target phenotype for all subsequent analyses.

For each genotype, Euclidean distance to the optimum phenotype was calculated as:

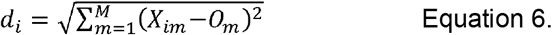

Where is the multidimensional Euclidean distance of genotype *i* to the optimal phenotype, *M* is the total number of traits included, represents the normalized value of trait *m* for genotype *i*, and represents the coordinate value of the optimum integrated phenotype centroid for trait *m* (the mean profile of the highest-yielding target cluster under drought stress).

### Quantification of plasticity trajectories

To determine whether plastic responses moved genotypes toward or away from the optimum phenotype, Euclidean distances were calculated under both well-watered and drought conditions. The change in distance between environments was then used to classify plasticity trajectories. Genotypes were classified into four response categories: (i) non-plastic, (ii) plastic away from the optimum, where drought responses increased the distance to the optimum phenotype, (iii) plastic toward the optimum, where drought responses reduced the distance to the optimum phenotype but did not reach the optimum centroid, and (iv) achieving the optimum phenotype, where drought responses minimized the distance to the optimum centroid. Differences in yield under drought and yield stability among plasticity trajectory classes were evaluated using ANOVA followed by Tukey’s HSD tests (p < 0.05).

## Results

### Whole root system plasticity is not associated with yield stability

We first tested whether genotypes exhibiting plasticity in a greater number of root traits displayed improved yield stability under drought. Plastic trait count was quantified by summing the number of root traits classified as plastic across nine root anatomical and architectural phenotypes for each genotype in maize and barley, and for each of the three plasticity metrics. Across all three plasticity metrics, whole-root-system plasticity i.e. plastic trait count, was not significantly associated with yield stability (Figure 1). Most genotypes exhibited plasticity in fewer than three root traits simultaneously, regardless of species or plasticity metric. Notably, no genotype displayed significant plasticity across all nine measured root phenotypes. These results indicate that extensive plasticity across multiple root traits is uncommon and that the number of plastic root phenotypes is not predictive of yield stability under drought.

**Figure 1:**
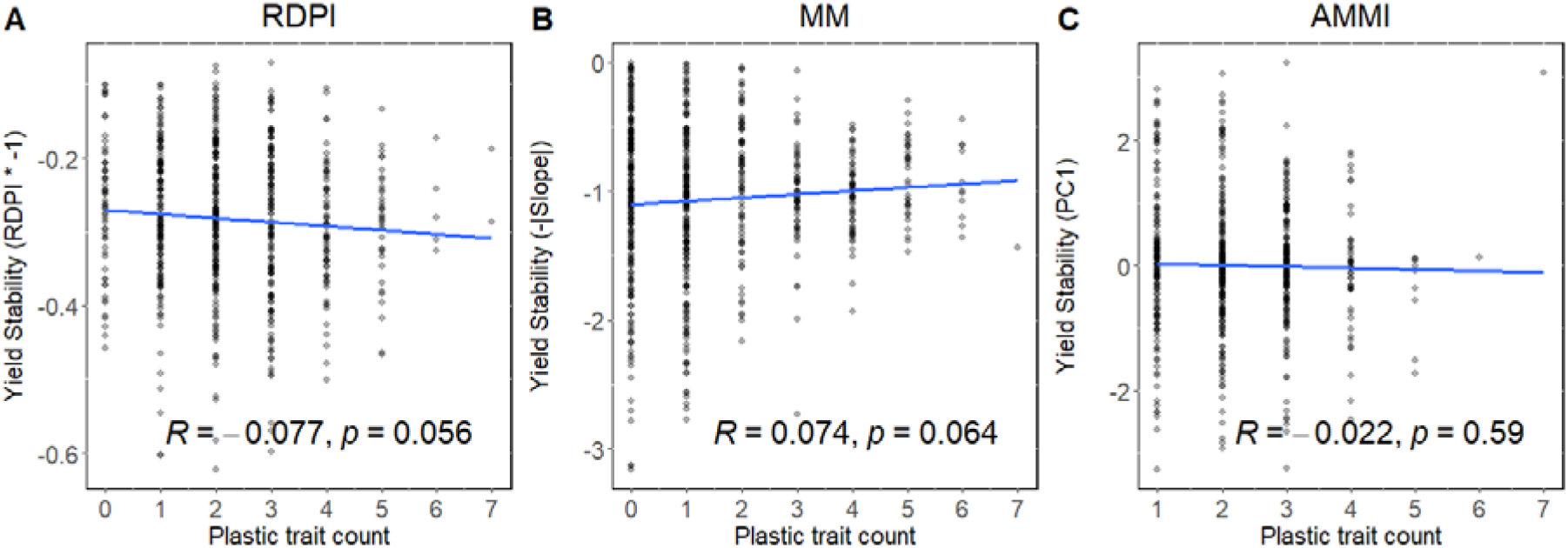
Relation between the plastic trait count and the yield stability for 621 genotypes (n=429 for maize and n=192 for barley). The root plasticity metrics include A) the Relative Distance Plasticity Index (RDPI), B) the slope of the Finlay–Wilkinson mixed model (MM), and C) the PC1 of the additive main effects and multiplicative interaction (AMMI) model. The units of the plastic trait count indicate the number of traits a specific genotype displayed plasticity out of 9 total traits (root angle, lateral root branching frequency, lateral root length, root cross-sectional area, root cortical aerenchyma, cortical cell size, cortical cell file number, number of metaxylem vessels, and median size of metaxylem vessels). A Pearson correlation test was performed to evaluate the association between the overall root plasticity and yield stability. The linear regression line is shown in blue. Each point represents a genotype and shows the plastic trait count and yield stability value derived from at least two replications per year and treatment.

### Greater cumulative plasticity does not improve yield or yield stability

To determine whether the magnitude of root plasticity influences crop performance, we calculated root system RDPI by summing absolute RDPI values across root phenotypes for each genotype. Unlike the plastic trait count, this metric incorporates both the number of plastic traits and the magnitude of the phenotypic response. In maize and barley, root system RDPI was not significantly associated with yield stability (Figure 2) or yield (Figure S6).

**Figure 2:**
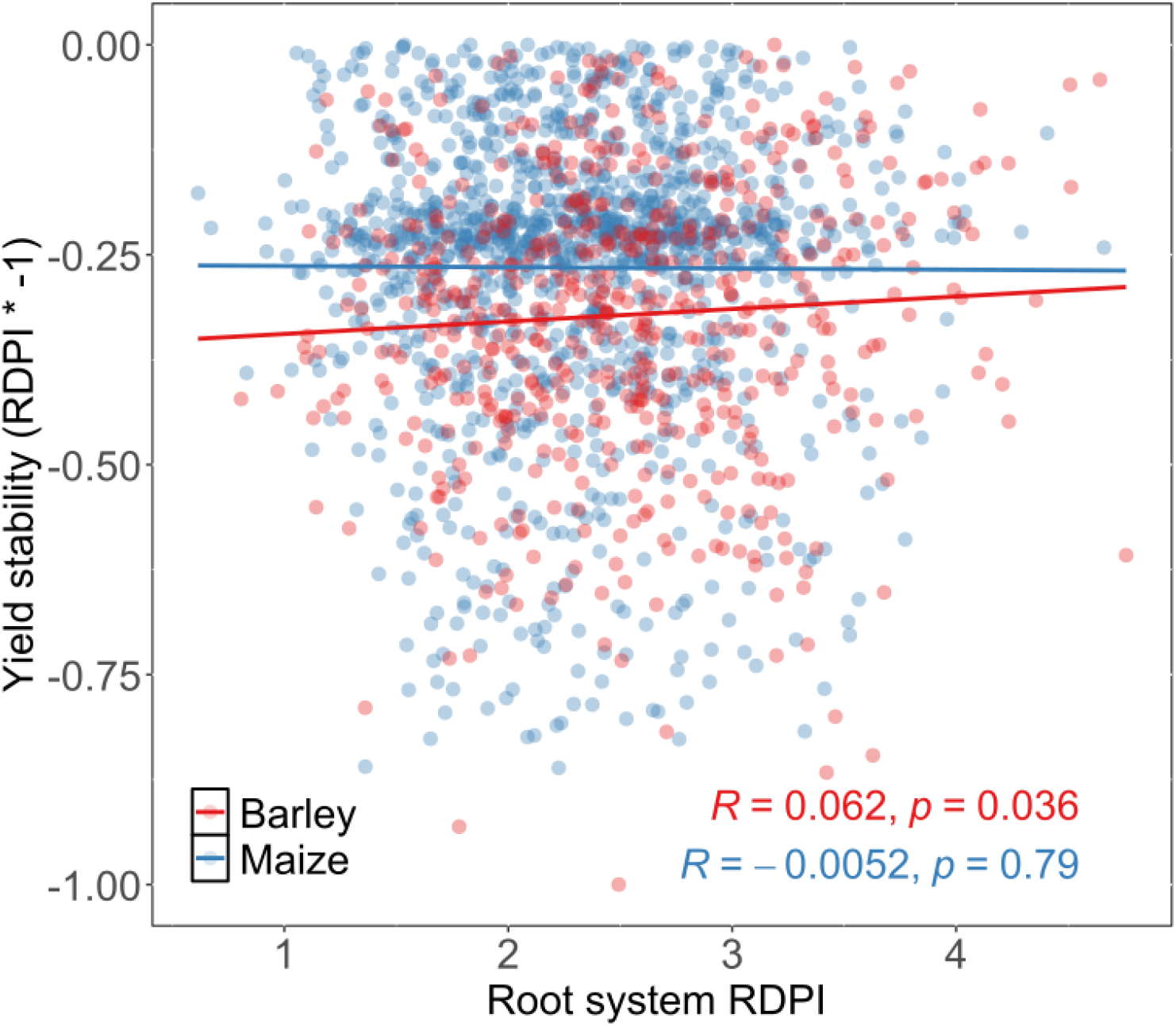
Relationship between root system RDPI (i.e. sum of all RDPI metrics of root traits) and yield stability. Each point represents the root system RDPI of different maize and barley genotypes, considering 9 root traits (root angle, lateral root branching frequency, lateral root length, root cross-sectional area, root cortical aerenchyma, cortical cell size, cortical cell file number, number of metaxylem vessels, and median size of metaxylem vessels). A Pearson correlation test was performed to evaluate the association. The linear regression line across each species is shown.

### Plasticity of individual root traits shows trait-dependent associations with yield stability

Because whole-root-system plasticity was not associated with yield stability, we next examined whether plasticity in specific root phenotypes contributed to crop performance. Correlations were calculated between yield stability and trait-specific plasticity estimates derived from RDPI, MM, and AMMI analyses. Associations between plasticity and yield stability were highly dependent on both the trait, species, and the plasticity metric used (Figure 3). In maize, plasticity in lateral root branching frequency (LRBF) was negatively correlated with yield stability. However, plasticity in lateral root branching frequency and lateral root length (LL) showed positive significant associations with yield under stress conditions for maize and barley, respectively, but not under well-watered conditions (Figure S7). These results indicate that plasticity in specific root phenotypes can be associated with improved crop performance, especially in stress conditions although the strength and consistency of these relationships depend on both species and the method used to quantify plasticity.

**Figure 3:**
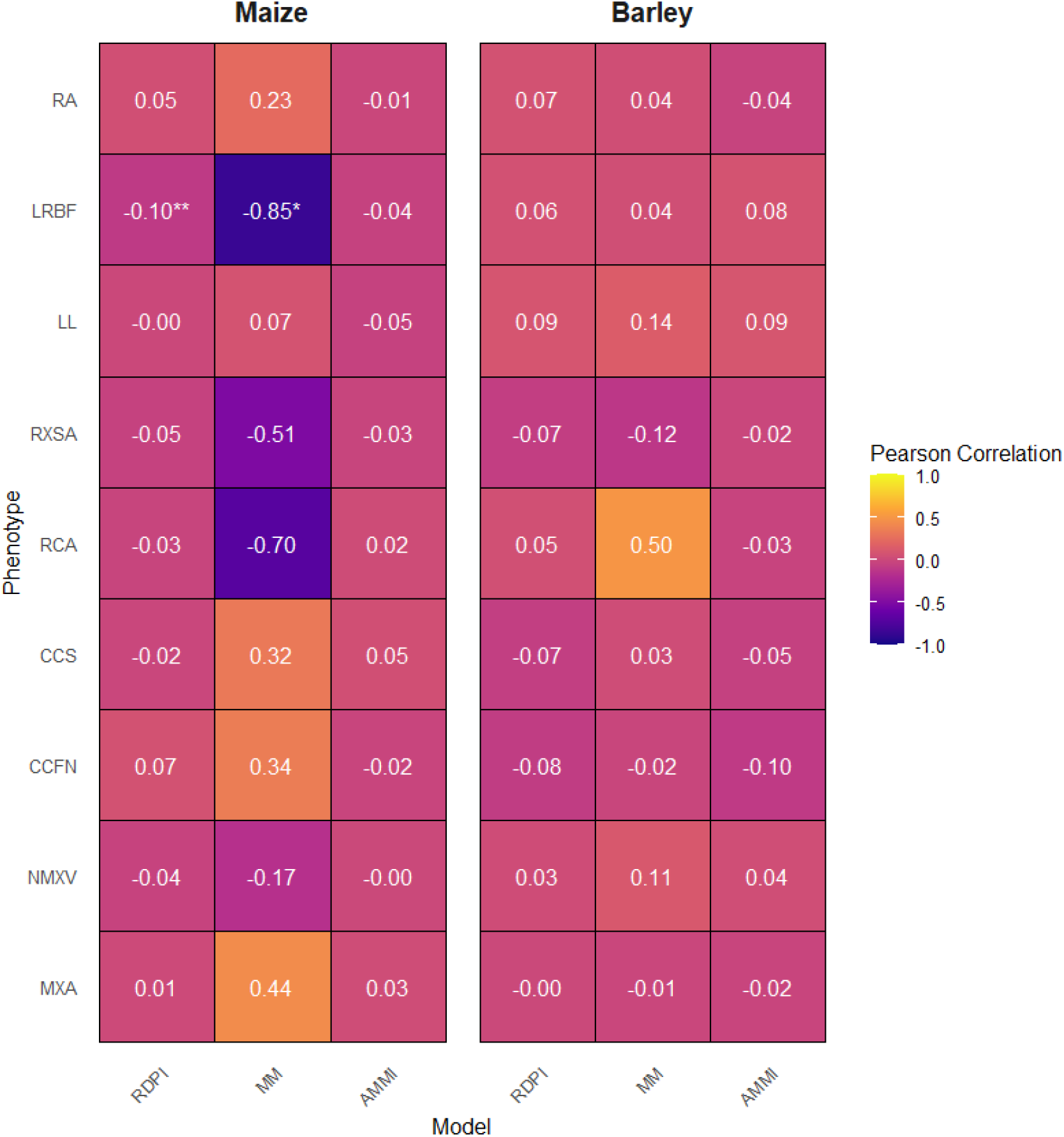
Correlation between the yield stability and the plasticity of individual root traits for maize and barley. Root trait plasticity was determined by Relative Distance Plasticity Index (absolute RDPI), slope of the Finlay–Wilkinson mixed model (MM), and additive main effects and multiplicative interaction (AMMI). Individual root traits include: root angle (RA), lateral

### Similar plastic responses occur in yield-stable and yield-unstable genotypes

To determine why plasticity in some root traits were associated with yield stability whereas others were not, genotypes were partitioned into yield-stable and yield-unstable groups using yield stability metrics derived from RDPI and MM analyses (Figure 4; Figures S5 and S8). Each group (yield-stable and yield-unstable) was then subgrouped using Partition Around Medoids (PAM) clustering, producing two stable groups (stable C1 and stable C2) and two unstable groups (unstable C1 and unstable C2), based on the coordinated plastic responses of nine root phenotypes.

**Figure 4:**
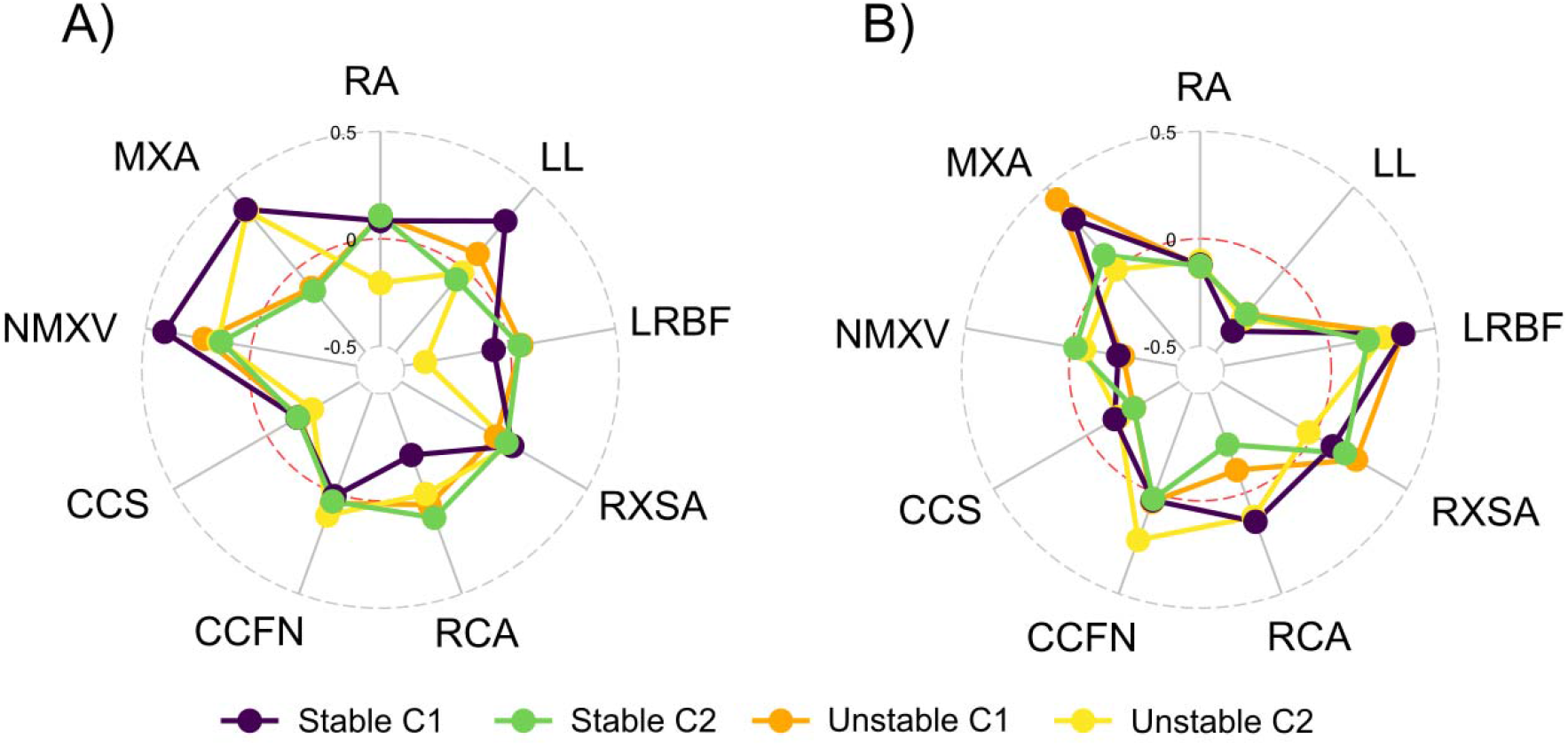
Root plasticity strategies for yield stable and unstable clusters for A) maize and B) barley. In each radar plot, we show four profiles corresponding to the root phenotypic plasticity of nine root traits. Profiles are divided into stable and unstable, and into clusters 1 and 2. ’Stable’ and ’Unstable’ correspond to the bulks obtained by filtering the stable genotypes from the unstable genotypes for yield (Figure S5). Numbers 1 and 2 correspond to the clusters of phenotypes (i.e. strategies) that represent the strategies within the yield stable and unstable groups. The plasticity value for each trait is the non-absolute RDPI. The red line indicates a non-absolute RDPI value of 0, meaning no plasticity. Positive values indicate a plastic response increasing the trait value, and negative values indicate a plastic response decreasing the trait value. The abbreviations for the phenotypes are: root angle (RA), lateral root branching frequency (LRBF), lateral root length (LL), root cross-sectional area (RXSA), root cortical aerenchyma (RCA), cortical cell size (CCS), cortical cell file number (CCFN), number of metaxylem vessels (NMXV), and median area of metaxylem vessels (MXA). root branching frequency (LRBF), lateral root length (LL), root cross-sectional area (RXSA), root cortical aerenchyma (RCA), cortical cell size (CCS), cortical cell file number (CCFN), number of metaxylem vessels (NMXV), and median area of metaxylem vessels (MXA). A Pearson correlation test was performed to evaluate the associations. Asterisks indicate statistical significance levels: *** for P < 0.01, ** for P < 0.05, and * for P < 0.1.

Surprisingly, several integrated plasticity profiles in response to drought were highly similar between yield stable and unstable groups (Figure 4). In maize, both yield stable C2 and unstable C1 exhibited increased metaxylem vessel number and reduced metaxylem vessel area, while yield stable C1 and unstable C1 shared increases in lateral root length and lateral root branching frequency. Likewise, in barley, yield stable and unstable clusters displayed similar reductions in cortical cell file number and increases in metaxylem vessel area. Rather than representing fundamentally different drought-response strategies, many yield stable and unstable groups occupied closely related regions of phenotype space and differed only in a small number of root phenotypes.

These findings suggest that yield stability is not determined by the presence or absence of a particular plastic response. Instead, drought performance appears to depend on how those responses reconfigure a genotype within phenotype space. Yield-stable and yield-unstable groups often exhibited highly similar plastic shifts, but they began from different constitutive phenotypes. Therefore, the same magnitude of plasticity could move one genotype toward a favorable configuration while pushing another away from it. Consequently, the adaptive value of plasticity cannot be inferred from the magnitude of individual trait responses alone but must be considered in the context of the broader integrated phenotype in which those responses occur. These results suggest that drought adaptation is determined less by individual plastic responses and more by whether plasticity moves a genotype toward a favorable integrated phenotype.

### Identification of a drought-associated integrated phenotype

Because similar plastic responses occurred in both yield stable and unstable genotypes, with significant differences in plastic responses in only a few root traits, we next investigated whether drought adaptation was associated with plasticity oriented towards a specific integrated phenotype rather than general plastic responses. To define the target integrated phenotype to which plasticity would be measured from, an exhaustive evaluation of more than 1,500 trait combinations was performed using Partition Around Medoids clustering across maize (Table S4) and barley (Table S5) under drought conditions.

For maize, among all evaluated trait combinations (Table S4), flowering time, cortical cell file number, and median metaxylem vessel area provided the strongest separation of genotypes in phenotype space based on silhouette width (Table S6) and significant differences for yield among clusters (Table S6). Within this configuration, cluster 2 consistently outperformed all others under drought stress (Figure 5A). The centroid of this cluster was defined as the optimum phenotype, underpinned by these specific phenological and root anatomical traits, representing the target integrated phenotype associated with superior yield drought performance (Table 2). To quantify the proximity of individual genotypes to this optimum phenotype, Euclidean distances were calculated between each genotype and the centroid of the optimum cluster (Figure 6). For barley, among all trait combinations (Table S5), lateral root branching frequency, root cross-sectional area and cortical cell size provided the strongest cluster structure by greater silhouette width values and significant differences for yield under drought (Table S7). For the case of barley, cluster 1 showed the highest yield under stress when using this particular trait configuration (Figure 5B, Table 3). Cluster 1 was then considered as the optimal phenotype for barley and the Euclidian distance towards this cluster was calculated for further analysis. Genotypes varied substantially in their distance from the optimum phenotype, indicating considerable variation in the degree to which constitutive root phenotypes approached the target phenotype associated with superior drought performance.

**Figure 5:**
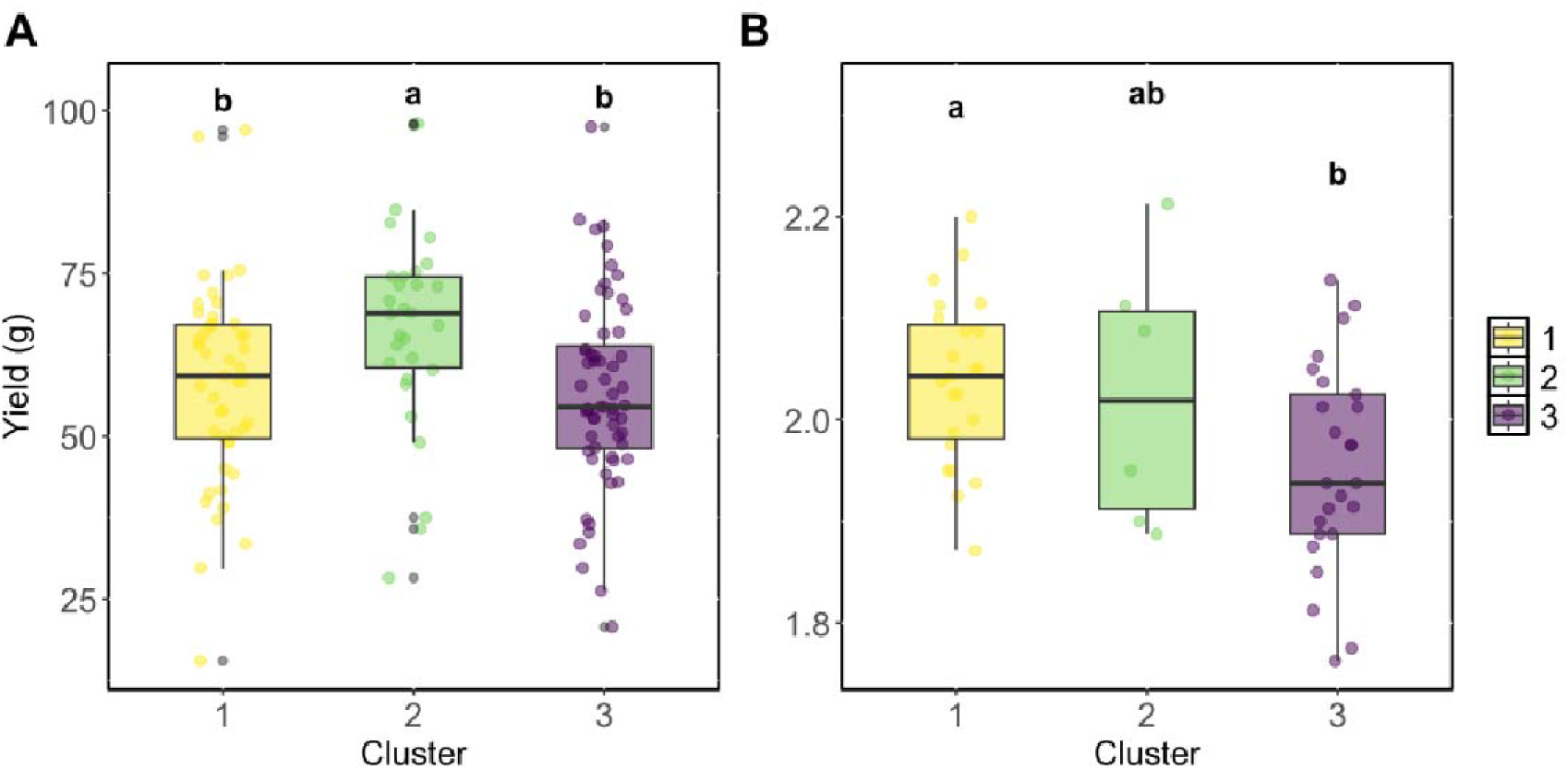
Yield differences among clusters found for A) maize and B) barley. Clusters were generated with the Medoids analysis using 3 different traits for maize and barley. The traits used for making the clusters and the average phenotypic state of each cluster can be found in Table 2 for maize and Table 3 for barley. The trait combination used in this analysis was selected due to its maximum silhouette values for strong cluster structure and significant yield differentiation of clusters under drought (Table S6 and Table S7). Different letters indicate significant statistical differences between means based on ANOVA and Tukey’s honest significant difference tests (P < 0.05).

**Figure 6:**
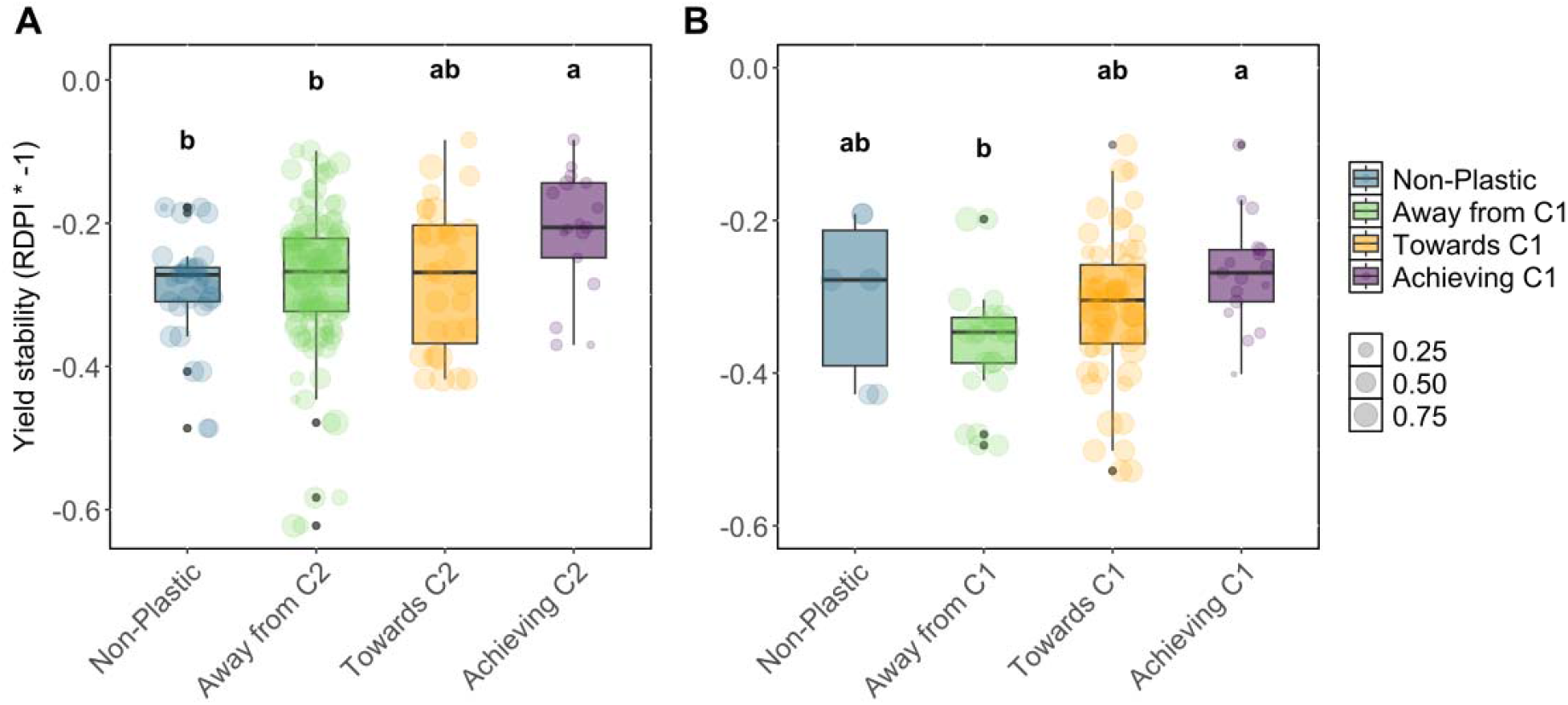
Yield stability differences among different classes of plasticity for A) maize and B) barley. Phenotypic plasticity was classified into three categories: "Away from C_" (plasticity driving the phenotype away from the optimal cluster), "Towards C_" (plasticity progressing toward optimal cluster but not fully reaching it), and "Achieving C_" (plasticity enabling complete achievement of the optimal cluster phenotype). For maize, cluster 2 was identified as the optimal cluster, and for barley, cluster 1. Distances to the optimum were therefore computed relative to the respective best cluster. Dot size denotes the Euclidian distance to optimal phenotype, with smaller dots indicating greater phenotypic similarity to the optimum. Different letters indicate significant statistical differences between means based on ANOVA and Tukey’s honest significant difference tests (P < 0.05).

### Plasticity toward the optimum phenotype is associated with improved drought performance

To determine whether the adaptive value of plasticity depended on its direction, genotypes were classified according to how their drought responses altered the distance to the optimum phenotype. Genotypes were categorized as moving away from the optimum phenotype, moving toward the optimum phenotype, or achieving the optimum phenotype following drought-induced plasticity (Figure 6). No significant differences among plasticity directions were observed under well-watered conditions (Figure S9A and S10A). Under drought stress, however, genotypes that achieved the optimum phenotype displayed significantly greater yield than genotypes exhibiting alternative plasticity trajectories (Figure S9B and S10B).

Yield stability also differed among plasticity classes (Figure 6). Genotypes whose plastic responses moved them toward or achieved the optimum phenotype exhibited significantly greater yield stability than genotypes whose responses increased the distance from the optimum phenotype. Furthermore, genotypes achieving the optimum phenotype were generally positioned closer to the optimum centroid than genotypes exhibiting alternative plasticity trajectories. Together, these results indicate that neither the number of plastic root traits nor the cumulative magnitude of root-system plasticity predicts drought performance. Instead, the adaptive value of plasticity is associated with whether phenotypic responses move a genotype toward a favorable integrated phenotype (Figure 7).

**Figure 7:**
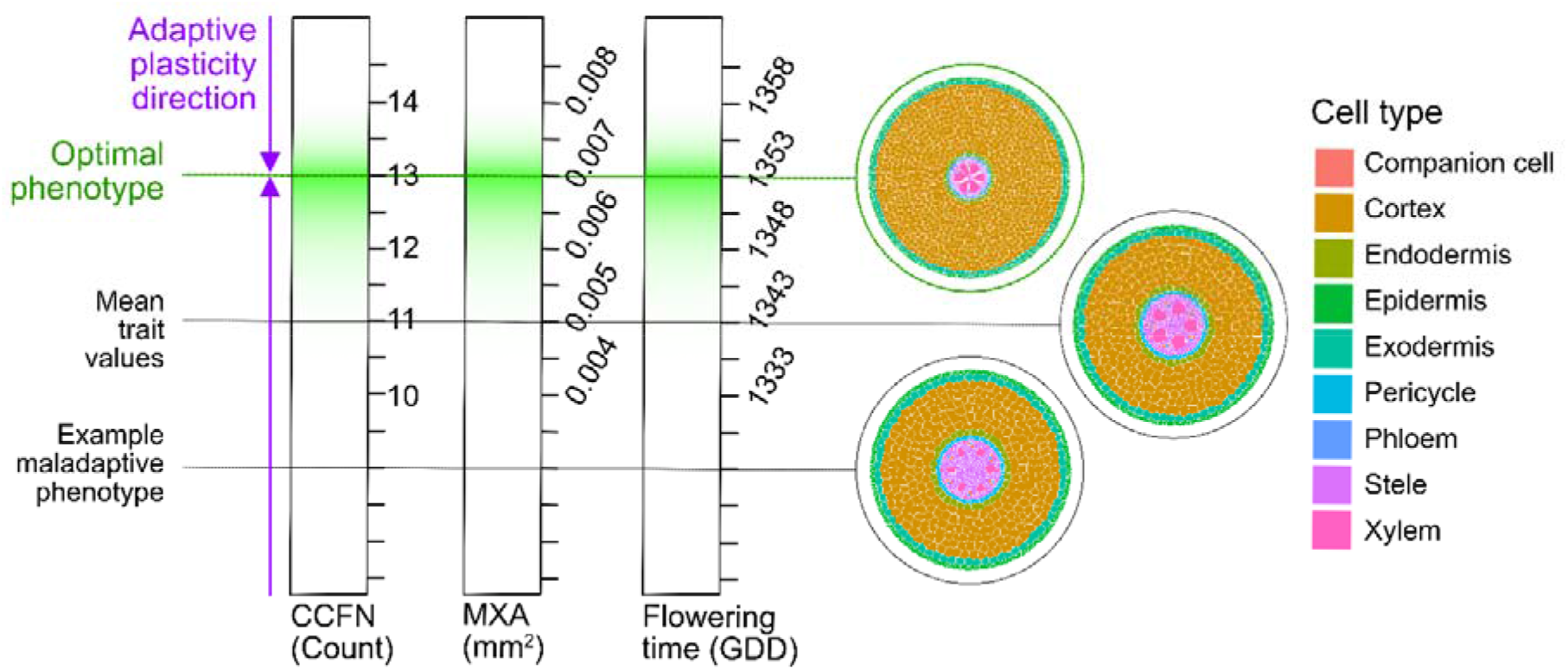
Schematic representation of plastic phenotypic trajectories in maize. The figure shows three vectors corresponding to the phenotypic space defined by cortical cell file number, metaxylem area, and flowering time. Green shading represents the phenotypic space of the optimal phenotype (i.e., cluster 2 in maize). A plastic response that moves the phenotype towards and achieves the optimal phenotype (purple arrow) has an adaptive value. Plastic responses that move the phenotype away from the optimal phenotype have a maladaptive value. Root anatomy sections representative of points of the phenotypic space were created using the online tool of GRANAR (Heymans et al., 2019).

## Discussion

Root plasticity is sometimes treated as a property of the root system as a whole. However, plasticity is not a monolithic trait of an entire organism. Rather, it is a modular property of individual roots, leaves, branches, and even specific tissues within those organs responding independently to environmental cues (De Kroon et al., 2005; Nielsen and Papaj 2022). Under this framework, plasticity emerges through the cumulative responses of numerous developmental modules rather than through coordinated whole-plant regulation. Consequently, there is little biological reason to expect that simply increasing the total amount of plasticity across the root system should consistently improve crop performance.

This perspective is strongly supported by our results. Across both maize and barley populations, most genotypes exhibited plasticity in only a small subset of the nine measured root phenotypes (Figure 1), with the majority displaying significant responses in fewer than three traits simultaneously. Notably, no genotype demonstrated significant plasticity across all nine traits (Figure 1). These observations indicate that environmental responses are genotype and trait-specific rather than coordinated across the entire root system. Consistent with this interpretation, neither the absolute number of plastic root traits nor the cumulative magnitude of root-system plasticity was associated with yield performance or yield stability under drought (Figure 1, 2, and S6).

### Is more plasticity necessarily better?

Root phenotypic plasticity has long been considered an attractive breeding target because it may enable crops to maintain performance across heterogeneous and unpredictable environments (Bradshaw 1965; Schneider, 2022). Our results do not support the assumption that greater plasticity is inherently adaptive. Across maize and barley, neither the number of plastic root traits nor the cumulative magnitude of root-system plasticity consistently predicted yield or yield stability (Figure 1 and 2). These findings suggest that selection is unlikely to favor plasticity itself. Rather, selection should favor specific plastic responses in specific traits that improve whole-plant performance under particular environmental conditions.

There are also biological reasons why extensive plasticity may fail to improve crop performance. Plastic responses require environmental sensing, signal transduction, and developmental reprogramming, all of which incur energetic and developmental costs (Auld et al., 2010). Furthermore, responses that improve one aspect of plant function may generate tradeoffs with other processes, particularly when resources are limiting (Schneider and Lynch 2020; Schneider, 2022). Consequently, greater phenotypic plasticity is not inherently advantageous if the resulting phenotype moves the plant away from an adaptive state. Under this perspective, the absence of a positive relationship between overall root-system plasticity and crop performance is not surprising.

### Drought adaptation emerges from coordinated trait combinations

Although plasticity in several individual root traits was associated with yield or yield stability, these relationships alone did not explain drought adaptation. For example, plasticity in lateral root branching frequency was associated with yield stability in maize (Figure 3), while plasticity in the same trait was also associated with yield under drought stress (Figure S7). However, there were no other distinct patterns between yield stability and yield under drought for most other root traits. More broadly, yield stable and unstable genotypes frequently exhibited remarkably similar plastic responses despite substantial differences in yield performance (Figure 4). Several plasticity profiles identified among yield stable genotypes were also observed among unstable genotypes, indicating that the same individual trait responses can occur in genotypes with contrasting drought outcomes.

These observations suggest that drought adaptation cannot be explained by individual root traits in isolation. Instead, the adaptive value of a plastic response depends on the phenotypic context in which it occurs. Root systems are composed of numerous interacting anatomical and architectural phenotypes that collectively influence resource acquisition, metabolic costs, and hydraulic function (Klein et al., 2020; Ajmera et al., 2022). Consequently, a change in a single trait may have different consequences depending on the configuration of the surrounding phenotype.

This finding has important implications for how plasticity is interpreted in crop physiology. Much of the literature evaluates plastic responses as isolated shifts in individual traits (e.g. Gao and Lynch, 2016; Zhan et al., 2015; Uga et al. 2013; Schneider et al 2022). Yet root systems consist of numerous interacting anatomical and architectural phenotypes that collectively determine resource acquisition, metabolic costs, and hydraulic function (Klein et al 2020; Ajmera et al 2022). Consequently, identical changes in a single trait (in terms of magnitude and direction) can produce very different final phenotypes depending on the initial phenotypic configuration. Under this framework, plasticity is better viewed as a trajectory through phenotype space rather than as a discrete change in trait values. The biological significance of a plastic response therefore depends not only on its magnitude but also on its starting position and final phenotype achieved.

The strong dependence of our results on the plasticity metric used further supports this interpretation. Significant relationships between plasticity and yield stability were most consistently identified using the Finlay–Wilkinson mixed model, whereas RDPI and AMMI often failed to detect the same trends (Figure 3). One explanation is that the Finlay– Wilkinson framework explicitly estimates genotype-specific reaction norms and allows confidence intervals to be used to distinguish consistent environmental responses from random variation. In contrast, RDPI and AMMI primarily quantify the magnitude of change without explicitly considering the repeatability or directional consistency of that response. While all three approaches provide useful information, our results suggest that identifying adaptive plasticity may require approaches that capture the direction and consistency of phenotypic trajectories rather than simply their magnitude.

### Does adaptive plasticity depend on movement toward an integrated phenotype?

If plasticity is fundamentally a trajectory through phenotype space, we must be able to define the destination or final state. Utilizing our feature discovery pipeline, we identified a highly specific target defined by a greater time to flowering, cortical cell file number, and median metaxylem vessel area for maize (Table 2, Figure 7) and lateral root branching frequency, root cross-sectional area and cortical cell size for barley (Table 3). Genotypes whose plasticity was oriented towards and achieving the optimal phenotype showed higher yield stability and yield (Figure 6, S9 and S10). Interestingly, more than 1,500 phenotypic combinations showed that adding additional traits to the model did not improve the cluster structure (Table S4 and S5). This indicates that a relatively small set of strategically important features is sufficient to capture the key physiological tradeoffs associated with drought adaptation.

The biological relevance of the identified phenotype is supported by the known functions of its component traits. Flowering time influences the temporal relationship between crop development and water availability and is frequently associated with drought escape strategies (Klein et al., 2020). Metaxylem vessel size influences hydraulic conductance and therefore affects the ability of roots to transport water from deeper soil layers to the shoot (Strock et al., 2026; Klein et al., 2024). Cortical cell file number affects root construction costs and may influence the metabolic investment required to build and maintain the root system (Chimungu et al., 2014; Lopez-Valdivia et al., 2023). Together, these traits in maize potentially integrate developmental timing, hydraulic function, and root metabolic costs, three processes that strongly influence drought adaptation. Similarly in barley, lateral root branching frequency determines soil foraging and regulates intra-plant competition (Zhan et al 2015), while root cross-sectional area and cortical cell size regulate hydraulic capacity and metabolic investment (Chimungu et al., 2014; Lopez-Valdivia et al., 2023), collectively forming an integrated barley root phenotype that enhances performance under water-limited conditions.

Most importantly, genotypes whose plastic responses moved them toward the optimum phenotype exhibited greater drought yield and yield stability than genotypes whose responses moved them away from it (Figure 6, S9, S10). Our trajectory mapping framework demonstrates that adaptive plasticity cannot be evaluated by measuring the sheer volume of phenotypic change. Instead, its value is entirely dependent on spatial orientation. A plastic response is only beneficial if the trajectory bridges the gap between the constitutive starting point and the optimal target. This interpretation is consistent with the argument of Sadras and Denison (2016) that optimization of individual traits in isolation is unlikely to maximize whole-plant performance because crop productivity emerges from interactions among multiple biological processes.

Our results also have implications for breeding. Historically, breeding programs have focused primarily on aboveground traits, with root traits receiving relatively little direct selection. When root traits have been targeted, the focus has typically been on individual traits in isolation, implicitly assuming that adaptive value resides within single traits (Richards and Passioura, 1989). Our results suggest that selecting for individual plastic responses may be insufficient because identical responses can have contrasting outcomes depending on the constitutive phenotype. Instead, breeding programs may benefit from identifying favorable integrated phenotypes and selecting genotypes capable of moving toward those phenotypic states under stress.

### Limitations and future directions

Several limitations should be considered when interpreting these results. First, all root phenotypes were measured at maturity. Consequently, transient developmental responses occurring earlier in the growing season may not have been captured. While several anatomical traits remain fixed following maturation, other traits such as lateral root emergence and root cortical aerenchyma can continue to change throughout development. Future studies incorporating repeated measurements through time will be necessary to determine how phenotypic trajectories evolve during plant ontogeny.

Second, root phenotypes explain only a portion of the variation in crop yield. Yield is influenced by numerous developmental, physiological, and environmental processes operating above and below ground. Consequently, even highly informative root traits are expected to explain only a fraction of total yield variation in diverse germplasm panels. Finally, the optimum phenotype identified in this study should not be interpreted as a universal drought ideotype. Different germplasm pools, cropping systems, and drought scenarios are likely to possess alternative adaptive phenotypic states. Rather than identifying a single universal optimum, the framework presented here provides a means of quantifying whether plastic responses move genotypes toward favorable integrated phenotypes within a given population and environment.

Our results demonstrate that whole-root-system plasticity is not a reliable predictor of yield or yield stability in maize and barley. Instead, the adaptive value of plasticity appears to depend on the direction of phenotypic change and whether that change moves a genotype toward a favorable integrated phenotype. These findings suggest that breeding for climate resilience should move beyond selecting for greater plasticity and instead focus on identifying phenotypic trajectories towards integrated phenotypes associated with improved performance under stress.

## Acknowledgements

We thank the IT department at Leibniz Institute of Plant Genetics & Crop Plant Research (IPK) for facilitating the computational resources and technical support required for this investigation.

## Funding

This study was supported by the Grains Research and Development Corporation ‘Root structure and function traits: Overcoming the root phenotyping bottleneck in cereals’ project (D.H.J.), the Leibniz Association (Resilient Roots to H.M.S and M.S.), and the European Union (ERC, 101162856, FATE) (H.M.S.). The views and opinions expressed are, however, those of the author(s) only and do not necessarily reflect those of the European Union or the European Research Council. Neither the European Union nor the granting authority can be held responsible.

### Declaration of Competing Interest

The authors declare that they have no known competing financial interests or personal relationships that could have appeared to influence the work reported in this paper.

*Declaration of generative AI and AI assisted technologies in the manuscript preparation process* During the preparation of this work, the author(s) used AI assisted tools for grammar checking and for troubleshooting in R code. After using these tools, the author(s) reviewed and edited all content and code outputs as needed and take full responsibility for the accuracy and integrity of the manuscript.

### Data availability

All data and scripts required to reproduce the analyses presented in this study are publicly available in the following repository: https://github.com/ilovaldivia/LopezValdivia2026_Root_plasticity *CRediT authorship contribution statement* **Ivan Lopez-Valdivia:** Writing – original draft, Writing – review & editing, Conceptualization, Formal analysis. **Akshay B. Tawale:** Writing – review & editing, Formal analysis. **Matías Schierenbeck:** Writing – review & editing, Formal analysis. **Daniele Sandoni:** Writing – review & editing, Formal analysis. **Dylan H. Jones:** Writing – review & editing. **Gwendolyn K. Kirschner:** Writing – review & editing. **Hannah M. Schneider:** Writing – original draft, Writing – review & editing, Conceptualization, Data curation, Project administration, Supervision.

---

**Figure S1:**
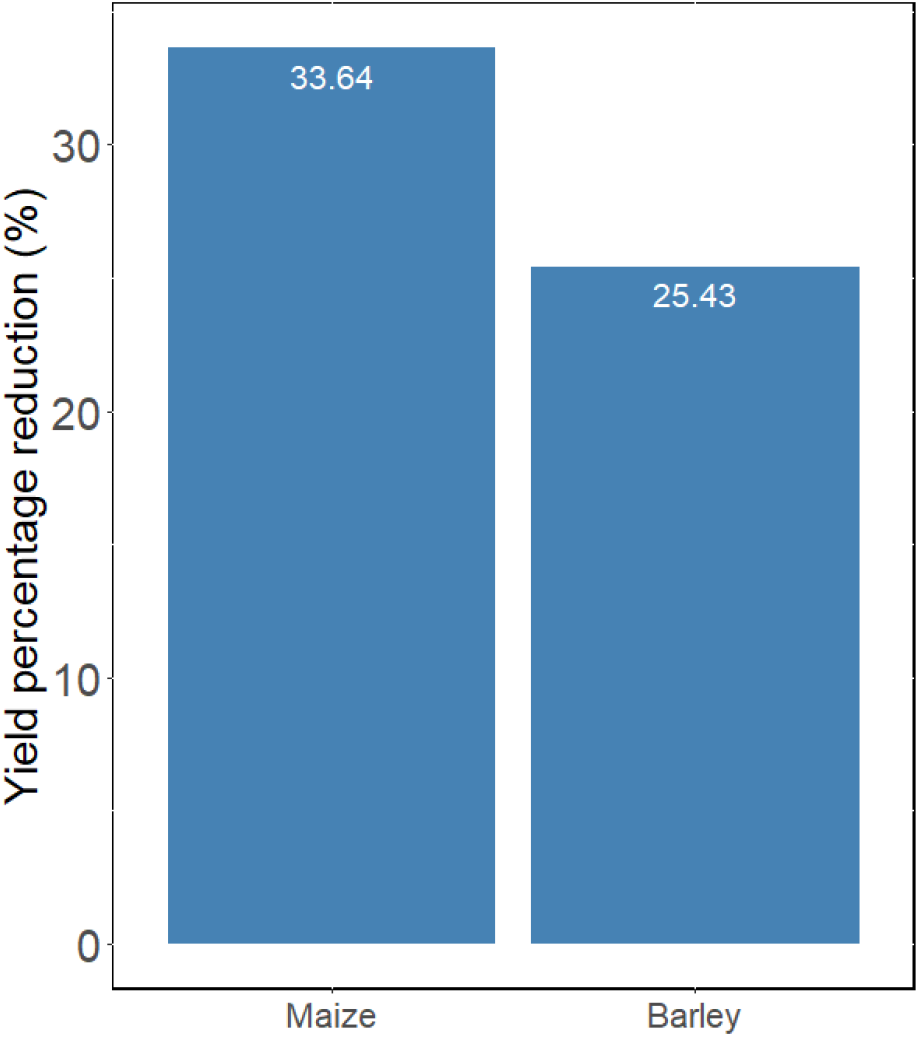
Yield reduction percentage under drought stress conditions for maize and barley. The bars represent the average percentage reduction in yield due to drought across 429 maize and 192 barley accessions.

**Figure S2:**
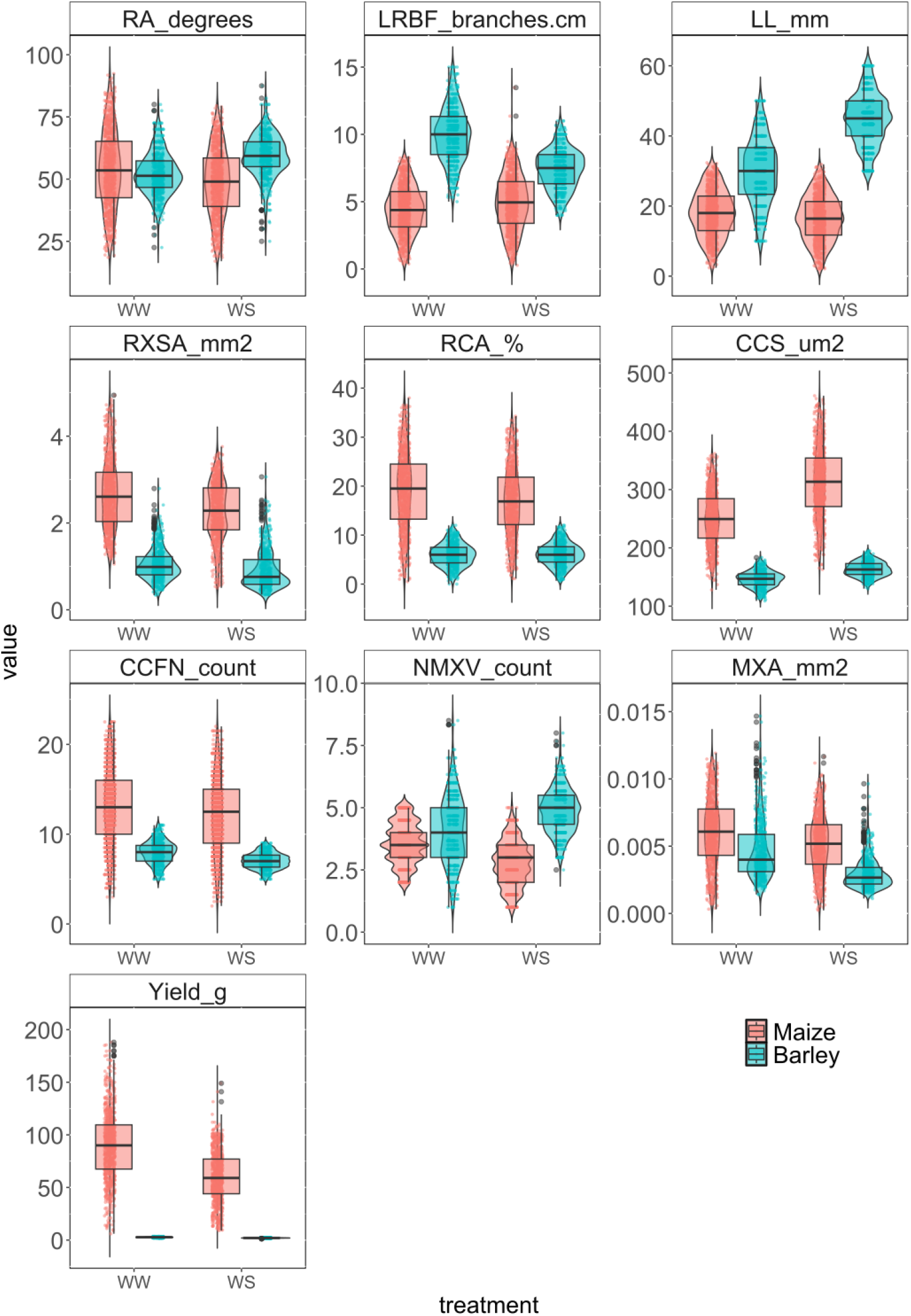
Root phenotypic values for 9 root traits across 2 species. Treatments are denoted by WW: well-watered, WS: water-stressed. Maize was grown in Arizona for 2 years (2015, 2016), barley was grown in Bonn for 3 years (2013, 2014, and 2015). The abbreviations for the phenotypes are: root angle (RA), lateral root branching frequency (LRBF), lateral root length (LL), root cross-sectional area (RXSA), root cortex area (RCA), cortical cell size (CCS), cortical cell file number (CCFN), number of metaxylem vessels (NMXV), and median area of metaxylem vessels (MXA).

**Figure S3:**
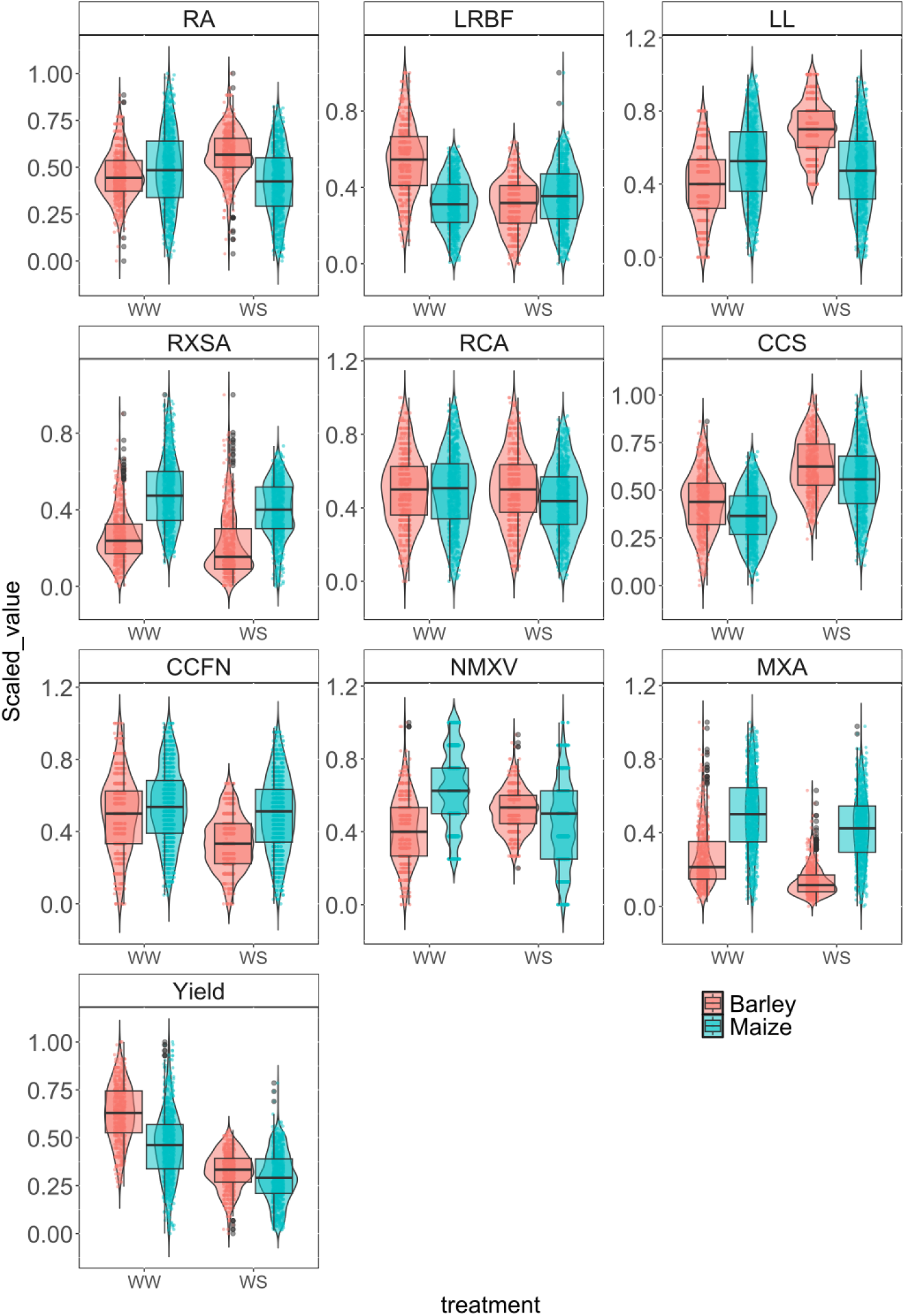
Normalized phenotypic data to a range from 0 to 1 using the min-max normalization method. Treatments are denoted by WW: well-watered, WS: water-stressed. Maize was grown in Arizona, USA for 2 years (2015, 2016), barley was grown in Bonn, Germany for 3 years (2013, 2014, and 2015). The abbreviations for the phenotypes are: root angle (RA), lateral root branching frequency (LRBF), lateral root length (LL), root cross- sectional area (RXSA), root cortex area (RCA), cortical cell size (CCS), cortical cell file number (CCFN), number of metaxylem vessels (NMXV), and median area of metaxylem vessels (MXA).

**Figure S4:**
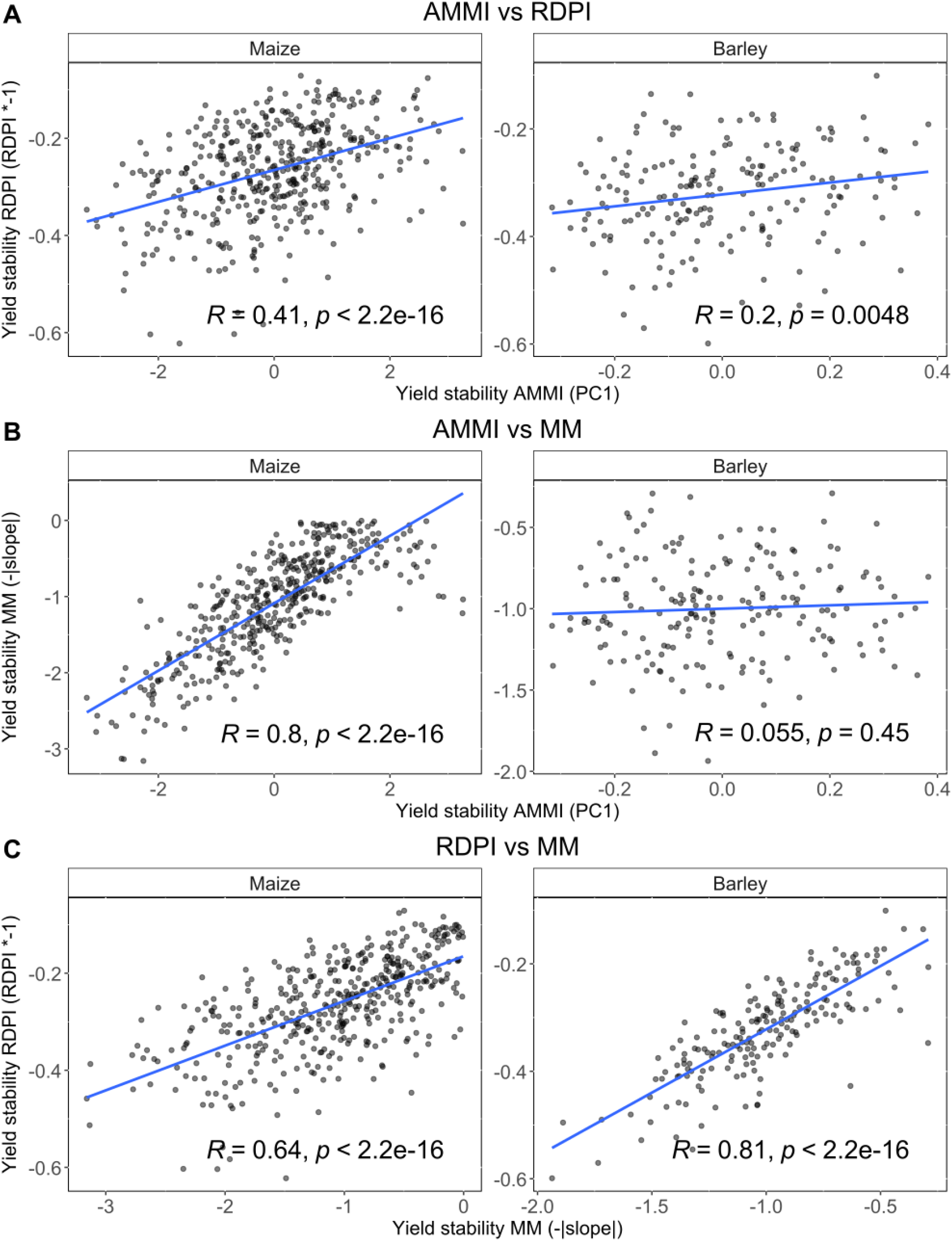
Associations between yield stability metrics for maize and barley. Stability metrics of yield include Relative Distance Plasticity Index (RDPI), slope of the Finlay–Wilkinson mixed model (MM), and additive main effects and multiplicative interaction (AMMI). A) shows the association between RDPI and AMMI; B) shows the association between MM and AMMI; and C) shows the association between MM and RDPI. R represents the Pearson correlation coefficient, accompanied by the P value. The linear regression line is shown in blue.

**Figure S5:**
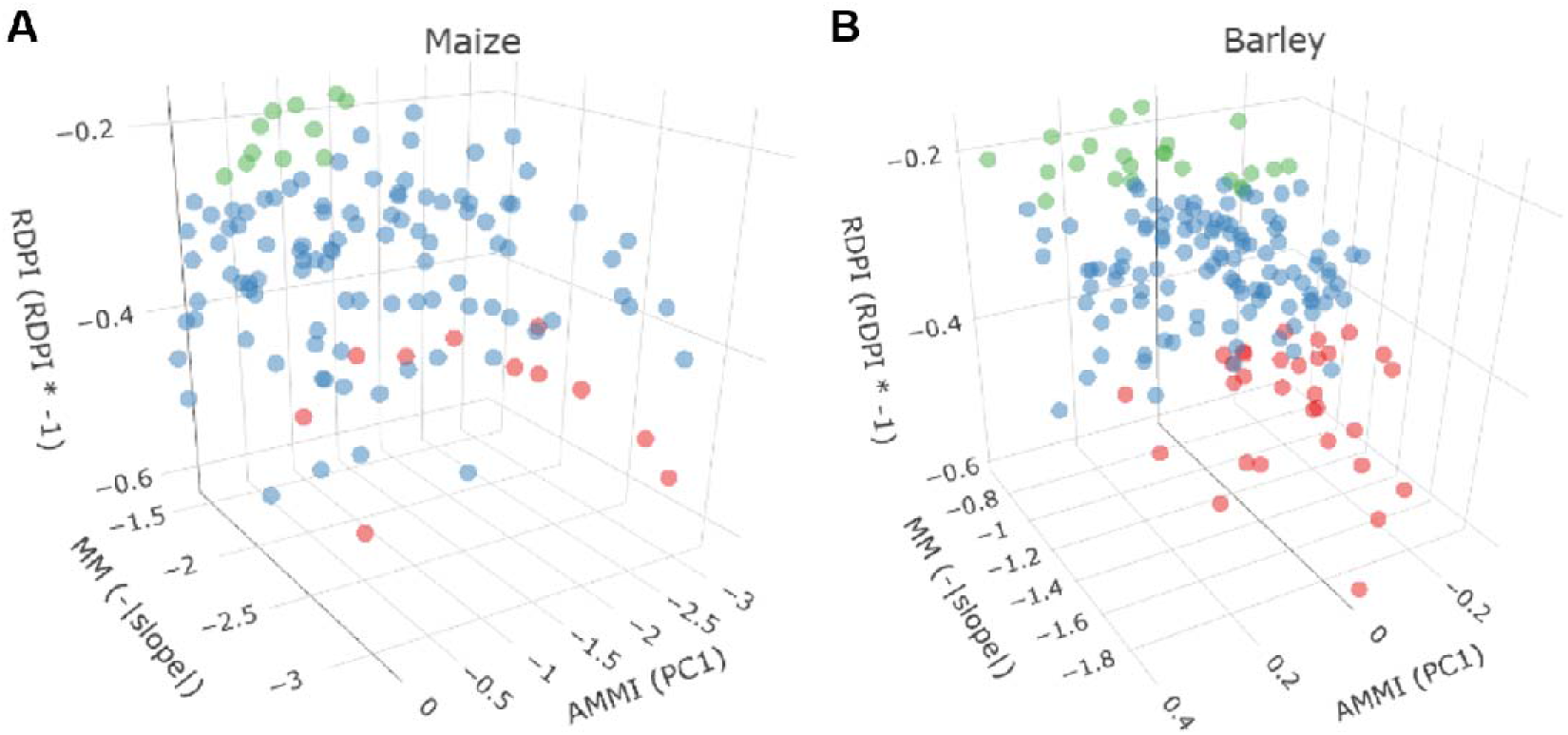
Yield stable and unstable groups based on plasticity metrics for a) maize and b) barley genotypes. Genotypes shown in green are stable for yield, while those in red are unstable. Genotypes in blue represent average yield stability. RDPI refers to the Relative Distance Plasticity Index; MM refers to the slope of the Finlay–Wilkinson model; and AMMI refers to the additive main effects and multiplicative interaction model.

**Figure S6:**
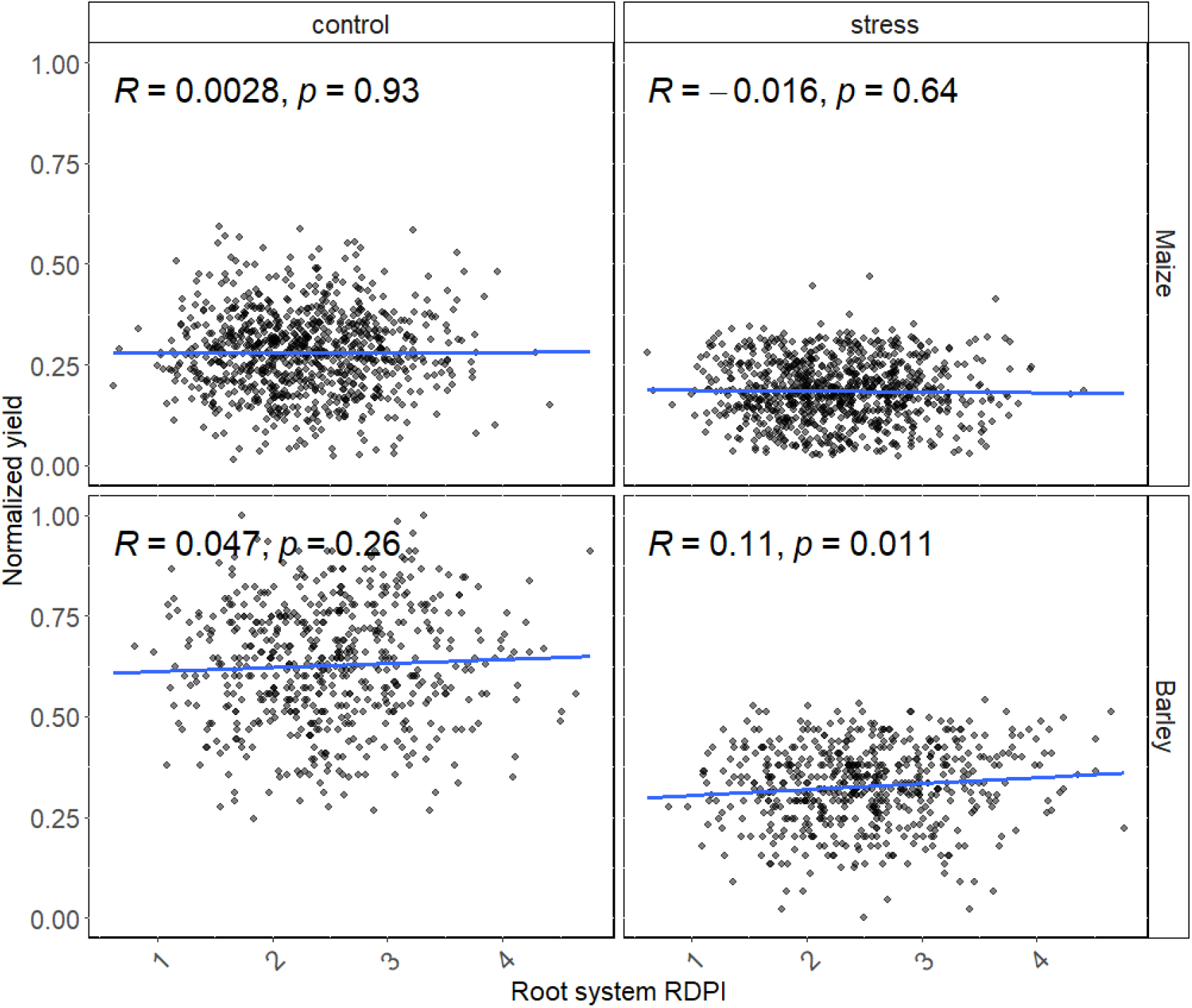
No association between normalized yield values and the total magnitude of plasticity (Root system RDPI), neither under control (WW), or water-stressed (WS) conditions. The genotypes included here are maize for two field seasons (2015, 2016) and barley grown for three field seasons (2013, 2014, and 2015). The linear regression line is shown in blue. Pearson correlation was performed and correlation coefficient is shown as R.

**Figure S7:**
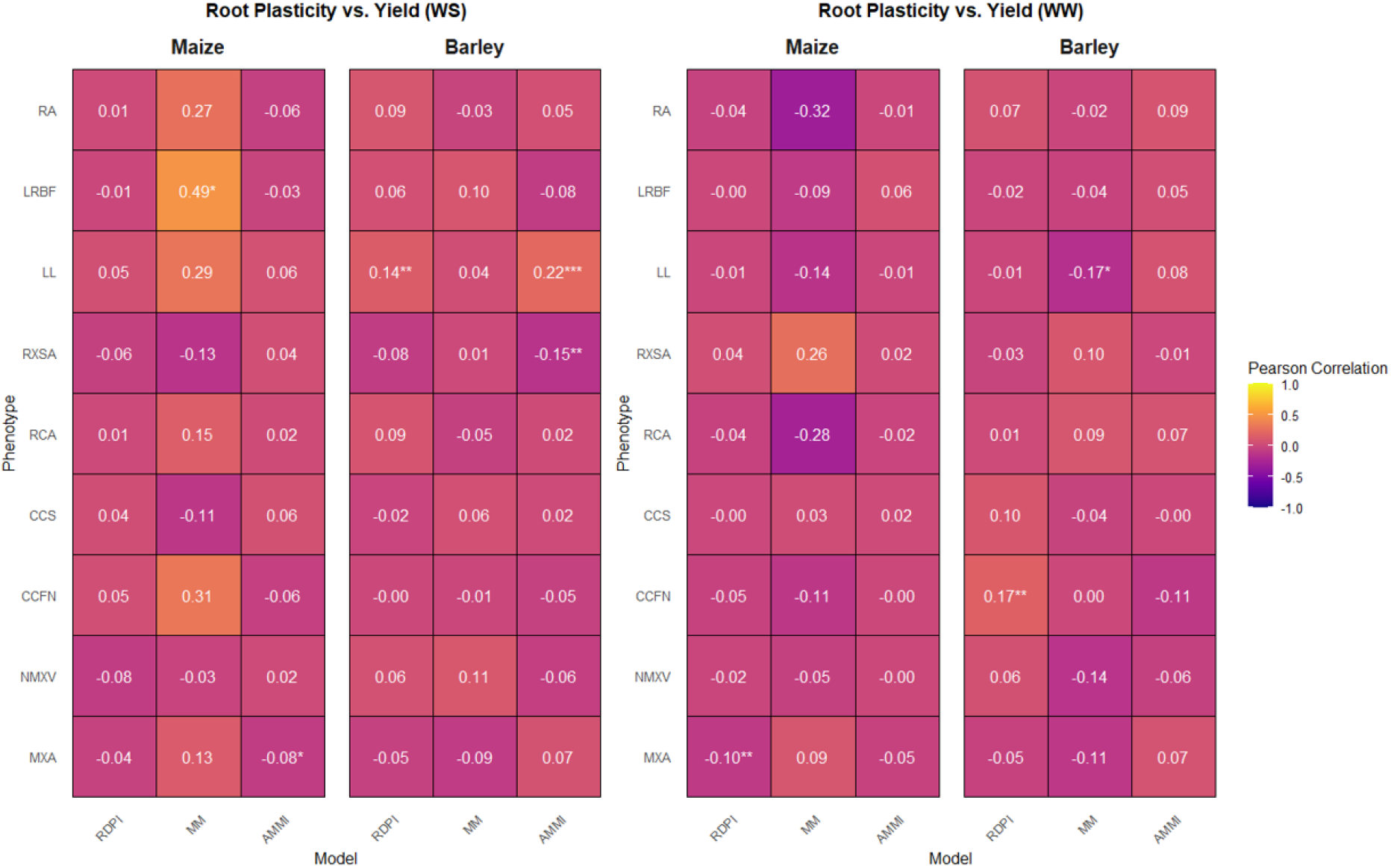
Correlation between scaled yield and individual root trait plasticity under control (WW) and drought stress (WS) conditions. Root trait plasticity was determined by Relative Distance Plasticity Index (RDPI), slope of the Finlay–Wilkinson mixed model (MM), and additive main effects and multiplicative interaction (AMMI). Individual root traits include: RA, root angle; LL, lateral root length; LRBF, lateral root branching frequency; RXSA, root cross-sectional area; RCA, root cortical aerenchyma; CCFN, cortical cell file number; CCS, cortical cell size; NMXV, number of metaxylem vessels; and MXA, median metaxylem vessel area. A Pearson correlation test was performed to evaluate the associations. Asterisks indicate statistical significance levels: *** for P < 0.01, ** for P < 0.05, and * for P < 0.1.

**Figure S8:**
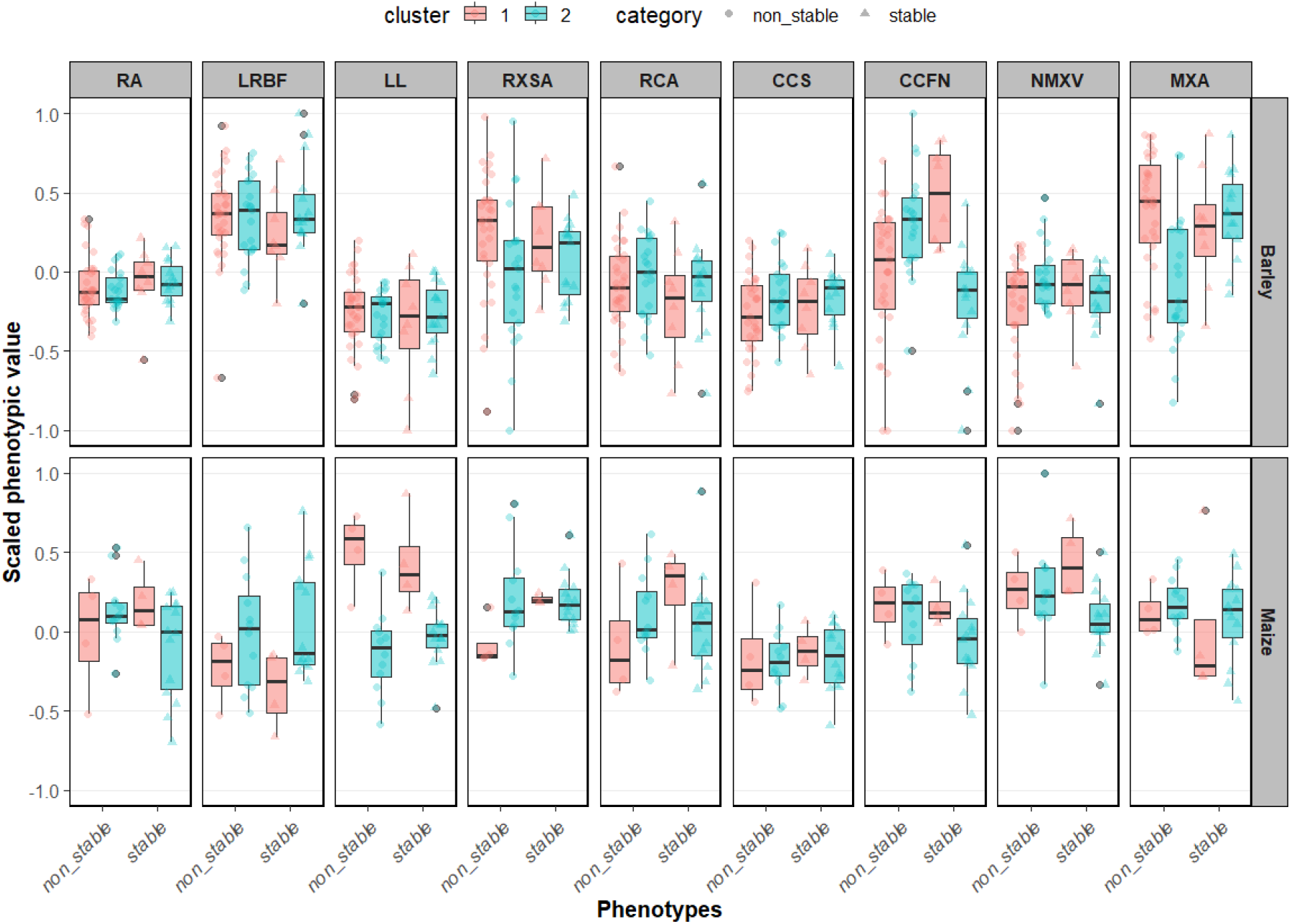
Scaled root phenotypes for yield stable and unstable groups in barley and maize under drought. Colors represent different clusters inside the stable and unstable groups. The y-axis shows the scaled root phenotypic values. Root phenotype abbreviations are as follows: RA, root angle; LL, lateral root length; LRBF, lateral root branching frequency; RXSA, root cross-sectional area; RCA, root cortical aerenchyma; CCFN, cortical cell file number; CCS, cortical cell size; NMXV, number of metaxylem vessels; and MXA, median metaxylem vessel area.

**Figure S9:**
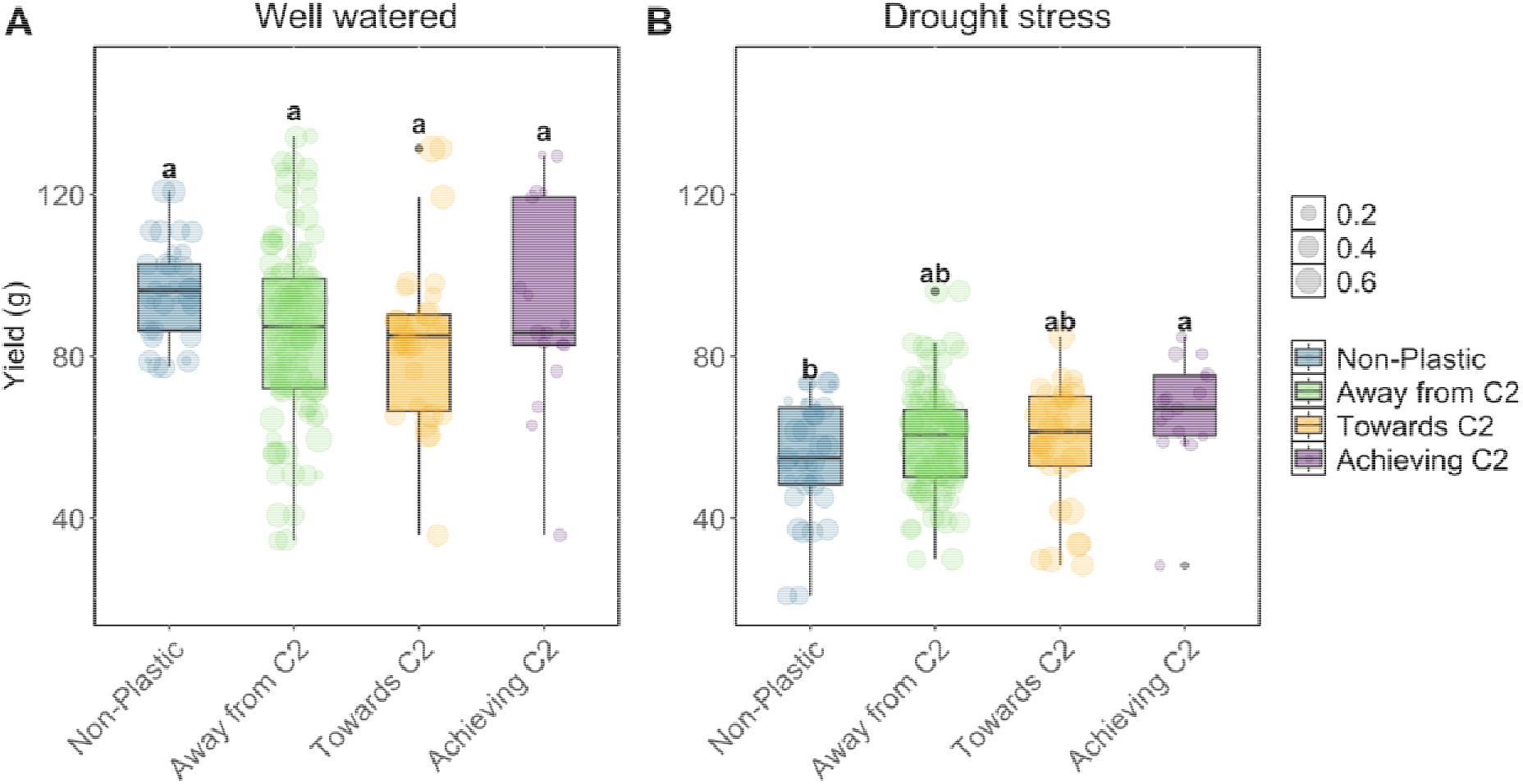
Different directions of plasticity and their association with yield for maize. Panels show the yield of the different categories of plasticity under A) control conditions and B) under drought stress conditions. Phenotypic plasticity was classified into three categories: "Away from C2" (plasticity driving the phenotype away from the optimal cluster 2), "Towards C2" (plasticity progressing toward optimal cluster 2 but not fully reaching it), and "Achieving C2" (plasticity enabling complete achievement of the optimal cluster 2 phenotype). Dot size denotes proximity to optimal phenotype 2, with smaller dots indicating greater phenotypic similarity to the optimum. Different letters indicate significant statistical differences between means based on ANOVA and Tukey’s honest significant difference tests (P < 0.05).

**Figure S10:**
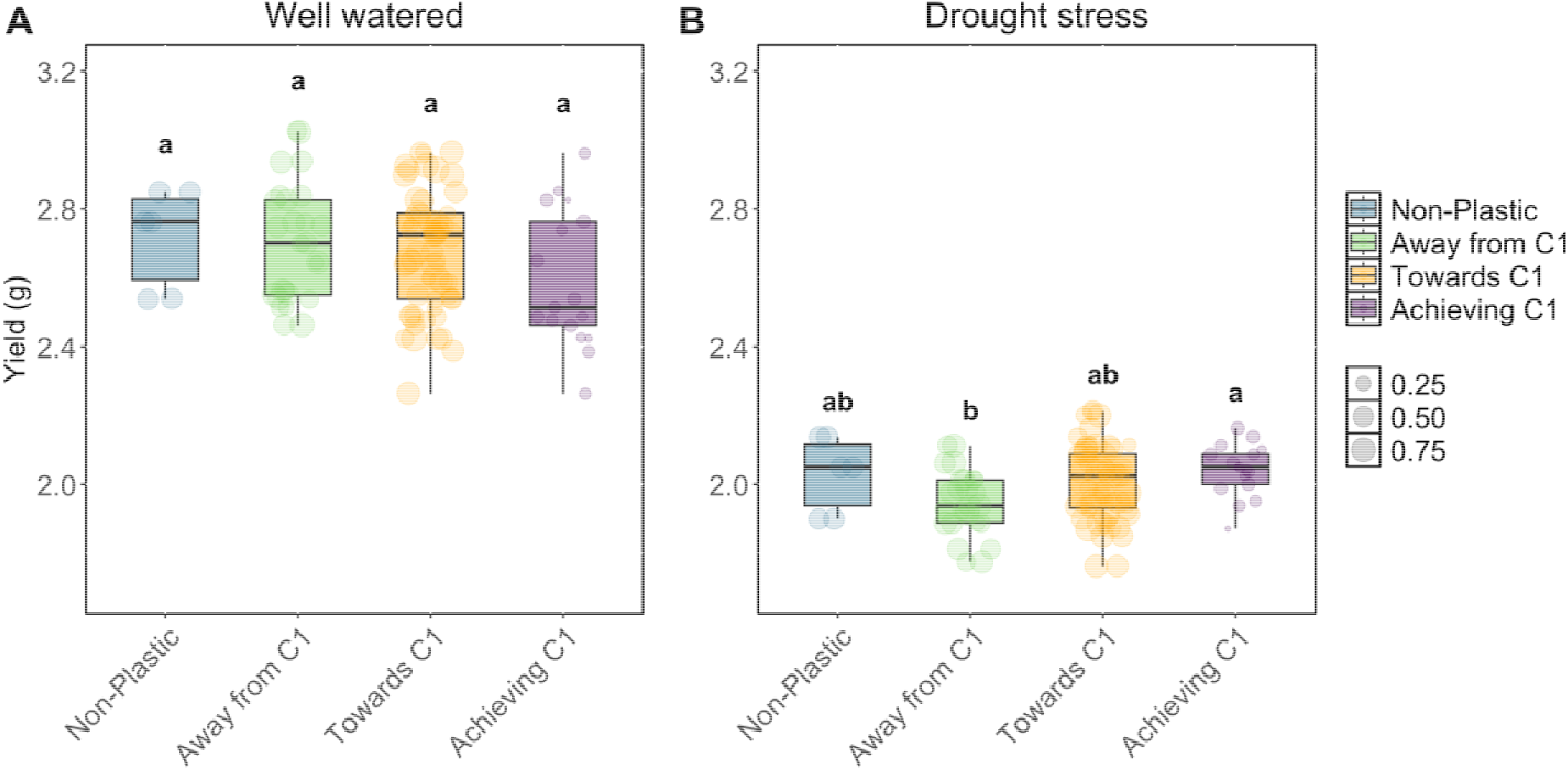
Different directions of plasticity and their association with yield for barley. Panels show the yield of the different categories of plasticity under A) control conditions and B) under drought stress conditions. Phenotypic plasticity was classified into three categories: "Away from C1" (plasticity driving the phenotype away from the optimal cluster 2), "Towards C1" (plasticity progressing toward optimal cluster 1 but not fully reaching it), and "Achieving C1" (plasticity enabling complete achievement of the optimal cluster 1 phenotype). Dot size denotes proximity to optimal phenotype 1, with smaller dots indicating greater phenotypic similarity to the optimum. Different letters indicate significant statistical differences between means based on ANOVA and Tukey’s honest significant difference tests (P < 0.05).

## References

1. Ajmera, I., Henry, A., Radanielson, A.M., Klein, S.P., Ianevski, A., Bennett, M.J., Band, L.R., Lynch, J.P. (2022). Integrated root phenotypes for improved rice performance under low nitrogen availability. Plant, Cell & Environment, 45, 805–822. 10.1111/pce.14284

2. Arenas, S., Djabali, Y., Rincent, R., Cubry, P., Martin, M.L., Blein-Nicolas, M., Laplaze, L., Schneider, H., Grondin, A. (2025). Modeling plant phenotypic plasticity and its underlying genetic architecture: a comparative study. Journal of Experimental Botany, 76, 2239–2251. 10.1093/jxb/eraf013

3. Auld, J.R., Agrawal, A.A., Relyea, R.A. (2010). Re-evaluating the costs and limits of adaptive phenotypic plasticity. Proceedings of the Royal Society B: Biological Sciences, 277, 503–511. 10.1098/rspb.2009.1355

4. Bao, Y., Aggarwal, P., Robbins, N.E., Sturrock, C.J., Thompson, M.C., Tan, H.Q., et al. (2014). Plant roots use a patterning mechanism to position lateral root branches toward available water. Proceedings of the National Academy of Sciences, 111, 9319–9324. 10.1073/pnas.1400966111

5. Bates, D., Mächler, M., Bolker, B., Walker, S. (2015). Fitting linear mixed-effects models using lme4. Journal of Statistical Software, 67, 1–48. 10.18637/jss.v067.i01

6. Bradshaw, A.D. (1965). Evolutionary significance of phenotypic plasticity in plants. Advances in Genetics, 13, 115–155.

7. Carrera, C.S., Savin, R., Slafer, G.A. (2024). Critical period for yield determination across grain crops. Trends in Plant Science, 29, 329–342. 10.1016/j.tplants.2023.08.012

8. Chimungu, J.G., Brown, K.M., Lynch, J.P. (2014). Reduced root cortical cell file number improves drought tolerance in maize. Plant Physiology, 166, 1943–1955. 10.1104/pp.114.249037

9. Correa, J., Postma, J.A., Watt, M., Wojciechowski, T. (2019). Soil compaction and the architectural plasticity of root systems. Journal of Experimental Botany, 70, 6019– 6034. 10.1093/jxb/erz383

10. De Kroon, H., Huber, H., Stuefer, J.F., van Groenendael, J.M. (2005). A modular concept of phenotypic plasticity in plants. New Phytologist, 166, 73–82. 10.1111/j.1469-8137.2004.01310.x

11. Ehdaie, B., Layne, A.P., Waines, J.G. (2012). Root system plasticity to drought influences grain yield in bread wheat. Euphytica, 186, 219–232. 10.1007/s10681-011-0585-9

12. Gao, Y., Lynch, J.P. (2016). Reduced crown root number improves water acquisition under water deficit stress in maize. Journal of Experimental Botany, 67, 4545–4557. 10.1093/jxb/erw243

13. Giehl, R.F.H., von Wirén, N. (2018). Hydropatterning—how roots test the waters. Science, 362, 1358–1359. 10.1126/science.aav9375

14. Hazman, M., Brown, K.M. (2018). Progressive drought alters architectural and anatomical traits of rice roots. Rice, 11, 1–14. 10.1186/s12284-018-0252-z

15. Heymans, A., Couvreur, V., LaRue, T., Paez-Garcia, A., Lobet, G. (2019). GRANAR, a computational tool to better understand the functional importance of monocotyledon root anatomy. Plant Physiology, 182, 707–720. 10.1104/pp.19.00707

16. Jennings, D.H., Trewavas, A.J. (eds.) (1986). Plasticity in Plants. Cambridge University Press.

17. Kadam, N.N., Tamilselvan, A., Lawas, L.M.F., Quinones, C., Bahuguna, R.N., Thomson, M.J., et al. (2017). Genetic control of plasticity in root morphology and anatomy of rice in response to water-deficit. Plant Physiology, 174, 2302–2315. 10.1104/pp.17.00500

18. Karlova, R., Boer, D., Hayes, S., Testerink, C. (2021). Root plasticity under abiotic stress. Plant Physiology, 187, 1057–1070. 10.1093/plphys/kiab392

19. Kassambara, A., Mundt, F. (2020). *factoextra: Extract and Visualize the Results of Multivariate Data Analyses*. R package version 1.0.7. 10.32614/CRAN.package.factoextra

20. King, J.G., Hadfield, J.D. (2019). The evolution of phenotypic plasticity when environments fluctuate in time and space. Evolution Letters, 3, 15–27. 10.1002/evl3.100

21. Klein, S.P., Kaeppler, S.M., Brown, K.M., Lynch, J.P. (2024). Integrating GWAS with a gene co-expression network best prioritizes candidate genes associated with root metaxylem phenes in maize. The Plant Genome, 17, e20323. 10.1002/tpg2.20323

22. Klein, S.P., Schneider, H.M., Perkins, A.C., Brown, K.M., Lynch, J.P. (2020). Multiple integrated root phenotypes are associated with improved drought tolerance. Plant Physiology, 183, 1011–1025. 10.1104/pp.20.00211

23. Knapp, S., van der Heijden, M.G.A. (2018). A global meta-analysis of yield stability in organic and conservation agriculture. Nature Communications, 9, 1–9. 10.1038/s41467-018-05956-1

24. Koehler, T., Kim, Y., Tung, S.Y., Heymans, A., Tyborski, N., Steiner, F., et al. (2025). Node of origin matters: comparative analysis of soil water limitation effects on nodal root anatomy in maize. Annals of Botany, 136, 1031–1046. 10.1093/aob/mcaf075

25. Lenth, R., Piaskowski, J. (2026). *emmeans: Estimated Marginal Means, aka Least- Squares Means*. R package version 2.0.4.

26. Li, N., Li, S., Zhao, B., Yuan, X., Zhu, J. (2026). Global agricultural drought crisis: synergistic impacts of climate change and human activities and their feedback mechanisms. Water, 18, 732. 10.3390/w18060732

27. Lopez-Valdivia, I., Rangarajan, H., Vallebueno-Estrada, M., Lynch, J.P. (2025). Broad environmental adaptation is associated with root anatomical phenotypes in maize landraces: an in silico study. Annals of Botany. 10.1093/aob/mcaf179

28. Lopez-Valdivia, I., Yang, X., Lynch, J.P. (2023). Large root cortical cells and reduced cortical cell files improve growth under suboptimal nitrogen in silico. Plant Physiology, 192, 2261–2275. 10.1093/plphys/kiad214

29. Lynch, J.P., Wojciechowski, T. (2015). Opportunities and challenges in the subsoil: pathways to deeper rooted crops. Journal of Experimental Botany, 66, 2199–2210. 10.1093/jxb/eru508

30. Maechler, M., Rousseeuw, P., Struyf, A., Hubert, M., Hornik, K. (2013). *cluster: Cluster Analysis Basics and Extensions*. R package.

31. Malika Charrad, N., Ghazzali, V., Boiteau, V., Niknafs, A. (2014). NbClust: An R package for determining the relevant number of clusters in a data set. Journal of Statistical Software, 61, 1–36. https://www.jstatsoft.org/v61/i06/

32. Mendiburu, F. (2023). *agricolae: Statistical Procedures for Agricultural Research*. R package version 1.3–7.

33. Mondal, K., da Costa, M.V.J., Sivarajan, S.R., Dey, N. (2026). Integrative plant responses to combined drought and heat stress: implications for climate-resilient crop development. Current Plant Biology, 48, 100625. 10.1016/j.cpb.2026.100625

34. Nielsen, M.E., Papaj, D.R. (2022). Why study plasticity in multiple traits? New hypotheses for how phenotypically plastic traits interact during development and selection. Evolution, 76, 858–869. 10.1111/evo.14464

35. Nijhout, H.F. (2003). Development and evolution of adaptive polyphenisms. Evolution & Development, 5, 9–18. 10.1046/j.1525-142x.2003.03003.x

36. Niones, J.M., Suralta, R.R., Inukai, Y., Yamauchi, A. (2012). Field evaluation on functional roles of root plastic responses on dry matter production and grain yield of rice under cycles of transient soil moisture stresses using chromosome segment substitution lines. Plant and Soil, 359, 107–120. 10.1007/s11104-012-1178-7

37. Niones, J.M., Suralta, R.R., Inukai, Y., Yamauchi, A. (2013). Roles of root aerenchyma development and its associated QTL in dry matter production under transient moisture stress in rice. Plant Production Science, 16, 205–216. 10.1626/pps.16.205

38. Oyiga, B.C., Palczak, J., Wojciechowski, T., Lynch, J.P., Naz, A.A., Léon, J., Ballvora, A. (2020). Genetic components of root architecture and anatomy adjustments to water-deficit stress in spring barley. *Plant*, Cell & Environment, 43, 692–711. 10.1111/pce.13683

39. Prince, S.J., Murphy, M., Mutava, R.N., Durnell, L.A., Valliyodan, B., Shannon, J.G., et al. (2017). Root xylem plasticity to improve water use and yield in water-stressed soybean. Journal of Experimental Botany, 66, 2027–2036. 10.1093/jxb/erw472

40. Richards, R.A., Passioura, J.B. (1989). A breeding program to reduce the diameter of the major xylem vessel in the seminal roots of wheat and its effect on grain yield in rain-fed environments. Australian Journal of Agricultural Research, 40, 943–950.

41. Sadras, V., Welsh, M., Sznajder, B., Hayes, J., Reynolds, M., Taylor, J. (2026). Rethinking yield stability through phenotypic plasticity and its link to modern statistical methods. *preprint*. 10.64898/2026.01.07.698075

42. Sandhu, N., Raman, K.A., Torres, R.O., Audebert, A., Dardou, A., Kumar, A., et al. (2016). Rice root architectural plasticity traits and genetic regions for adaptability to variable cultivation and stress conditions. Plant Physiology, 171, 2562–2576. 10.1104/pp.16.00705

43. Schneider, H.M. (2022). Characterization, costs, cues and future perspectives of phenotypic plasticity. Annals of Botany, 130, 131–148. 10.1093/aob/mcac087

44. Schneider, H.M., Klein, S.P., Hanlon, M.T., Kaeppler, S., Brown, K.M., Lynch, J.P. (2020b). Genetic control of root anatomical plasticity in maize. The Plant Genome, 13, e20003. 10.1002/tpg2.20003

45. Schneider, H.M., Klein, S.P., Hanlon, M.T., Nord, E.A., Kaeppler, S., Brown, K.M., Warry, A., Bhosale, R., Lynch, J.P., Dodd, I. (2020a). Genetic control of root architectural plasticity in maize. Journal of Experimental Botany, 71, 3185–3197. 10.1093/jxb/eraa084

46. Schneider, H.M., Lor, V.S.N., Hanlon, M.T., Perkins, A., Kaeppler, S.M., Borkar, A.N., Bhosale, R., Zhang, X., Rodriguez, J., Bucksch, A., Bennett, M.J. (2022). Root angle in maize influences nitrogen capture and is regulated by calcineurin B-like protein– interacting serine/threonine-protein kinase 15 (ZmCIPK15). Plant, Cell & Environment, 45, 837–853. 10.1111/pce.14135

47. Schneider, H.M., Lynch, J.P. (2020). Should root plasticity be a crop breeding target? Frontiers in Plant Science, 11, 546. 10.3389/fpls.2020.00546

48. Siangliw, J.L., Thunnom, B., Natividad, M.A., Quintana, M.R., Chebotarov, D., McNally, K.L., Lynch, J.P., Brown, K.M., Henry, A. (2022). Response of Southeast Asian rice root architecture and anatomy phenotypes to drought stress. Frontiers in Plant Science, 13, 1008954. 10.3389/fpls.2022.1008954

49. Strock, C.F., DePew, C.L., Sidhu, J.S., Xu, T., Lynch, J.P. (2026). A pleiotropic “stretch” phenotype is associated with metaxylem vessel element length, axial hydraulic conductance, root elongation, water utilization, and drought adaptation in maize. Crop Science, 66, 70287. 10.1002/csc2.70287

50. Strock, C.F., Schneider, H.M., Galindo-Castañeda, T., Hall, B.T., van Gansbeke, B., Mather, D.E., Roth, M.G., Chilvers, M.I., Guo, X., Brown, K.M., Lynch, J.P., Gifford, M. (2019). Laser ablation tomography for visualization of root colonization by edaphic organisms. Journal of Experimental Botany, 70, 5327–5342. 10.1093/jxb/erz271

51. Sultan, S.E. (1995). Phenotypic plasticity and plant adaptation. Acta Botanica Neerlandica, 44, 363–385.

52. Tebaldi, C., Lobell, D.B. (2008). Towards probabilistic projections of climate change impacts on global crop yields. Geophysical Research Letters, 35, L08803. 10.1029/2008GL033423

53. Thingujam, D., Gouli, S., Cooray, S.P., Chandran, K.B., Givens, S.B., Gandhimeyyan, R.V., Tan, Z., Wang, Y., Patam, K., Greer, S.A., Acharya, R., Moseley, D.O., Osman, N., Zhang, X., Brooker, M.E., Tagert, M.L., Schafer, M.J., Jeong, C., Hoffseth, K.F., Mukhtar, M.S. (2025). Climate-resilient crops: integrating AI, multi-omics, and advanced phenotyping to address global agricultural and societal challenges. Plants, 14, 2699. 10.3390/plants14172699

54. Tollenaar, M., Lee, E.A. (2002). Yield potential, yield stability and stress tolerance in maize. Field Crops Research, 75, 161–169.

55. Trachsel, S., Kaeppler, S.M., Brown, K.M., Lynch, J.P. (2011). Shovelomics: high throughput phenotyping of maize (Zea mays L.) root architecture in the field. Plant and Soil, 341, 75–87. 10.1007/s11104-010-0623-8

56. Uga, Y., Sugimoto, K., Ogawa, S., Rane, J., Ishitani, M., Hara, N., Kitomi, Y., Inukai, Y., Ono, K., Kanno, N., Inoue, H., Takehisa, H., Motoyama, R., Nagamura, Y., Wu, J., Matsumoto, T., Takai, T., Okuno, K., Yano, M. (2013). Control of root system architecture by DEEPER ROOTING 1 increases rice yield under drought conditions. Nature Genetics, 45, 1097–1102. 10.1038/ng.2725

57. Wang, T.-C., Moritz, A., Mabrouk, M., Villar Alegría, E., Arinalp, B., Ganji, E., Förter, L., Wittkop, B., Herzog, E., Snowdon, R.J., Stahl, A., Chen, T.-W. (2026). Plasticity of source–sink dynamics contributes to wheat yield stability. Nature Communications, 17, 3781. 10.1038/s41467-026-72330-x

58. Wasson, A.P., Richards, R.A., Chatrath, R., Misra, S.C., Prasad, S.V.S., Rebetzke, G.J., et al. (2012). Traits and selection strategies to improve root systems and water uptake in water-limited wheat crops. Journal of Experimental Botany, 63, 3485–3498. 10.1093/jxb/ers111

59. Wickham, H., François, R., Henry, L., Müller, K., Vaughan, D. (2023). dplyr: A Grammar of Data Manipulation. R package.

60. Xie, X., Quintana, M.R., Sandhu, N., Subedi, S.R., Zou, Y., Rutkoski, J.E., Henry, A. (2021). Establishment method affects rice root plasticity in response to drought and its relationship with grain yield stability. Journal of Experimental Botany, 72, 5208– 5220. 10.1093/jxb/erab214

61. Zhan, A., Schneider, H., Lynch, J.P. (2015). Reduced lateral root branching density improves drought tolerance in maize. Plant Physiology, 168, 1603–1615. 10.1104/pp.15.00187

